# *Legionella pneumophila* regulates host cell motility by targeting Phldb2 with a 14-3-3ζ-dependent protease effector

**DOI:** 10.1101/2021.09.06.459093

**Authors:** Lei Song, Jingjing Luo, Dan Huang, Yunhao Tan, Yao Liu, Kaiwen Yu, Yong Zhang, Xiaoyun Liu, Dan Li, Zhao-Qing Luo

## Abstract

The cytoskeleton network of eukaryotic cells is essential for diverse cellular processes, including vesicle trafficking, cell motility and immunity, thus is a common target for bacterial virulence factors. A number of effectors from the bacterial pathogen *Legionella pneumophila* have been shown to modulate the function of host actin cytoskeleton to construct the Legionella-containing vacuole (LCV) permissive for its intracellular replication. In this study, we identified the Dot/Icm effector Lem8 (Lpg1290) as a protease that interferes with host motility. We show that the protease activity of Lem8 is catalyzed by a Cys-His-Asp motif known to be associated with diverse biochemical activities. Intriguingly, we found that Lem8 interacts with the host regulatory protein 14-3-3ζ, which activates its protease activity. Furthermore, Lem8 undergoes self-cleavage in a process that requires 14-3-3ζ. We identified the PH domain-containing protein Phldb2 involved in cell migration as a target of Lem8 and demonstrate that Lem8 plays a role in the inhibition of host cell migration. Our results reveal a novel mechanism of inhibiting host cell motility by *L. pneumophila* for its virulence.

## Introduction

*Legionella pneumophila* is a Gram-negative intracellular bacterial pathogen ubiquitously found in freshwater habitats, where it replicates in a wide range of amoebae (Richards et al., 2013). It is believed that these natural hosts serve as the main replication niches for *L. pneumophila* in the environment and provide the primary evolutionary pressure for the acquisition and maintenance of virulence factors necessary for its intracellular lifecycle. Infection of humans by *L. pneumophila* occurs when susceptible individuals inhale aerosols generated from contaminated water, which introduces the bacterium to the lungs where it is phagocytosed by alveolar macrophages. Instead of being digested and cleared, internalized bacteria replicate within a membrane-bound compartment termed Legionella-containing vacuole (LCV), leading to the development of Legionnaires’ disease, a form of severe pneumonia (Cunha et al., 2016).

One feature associated with the LCV is its ability to evade fusion with the lysosomal network in the early phase (<8 h post-infection in mouse bone marrow-derived macrophages(BMDMs)) of its development and the quick acquisition of proteins of the endoplasmic reticulum (ER) origin (Kagan and Roy, 2002; Sturgill-Koszycki and Swanson, 2000; Swanson and Isberg, 1995). Biogenesis of the LCV requires the Dot/Icm type IV secretion system that injects more than 300 effector proteins into host cells (Qiu and Luo, 2017). These effectors function to modulate a wide cohort of host processes, including vesicle trafficking (Tan et al., 2011), protein synthesis (Shen et al., 2009), lipid metabolism (Gaspar and Machner, 2014), and autophagy (Choy et al., 2012) by diverse biochemical mechanisms. Coordinated activity of these effectors leads to the formation of the LCV which largely resembles the ER in its morphology and protein composition (Qiu and Luo, 2017).

The cytoskeleton of eukaryotic cells is composed of microfilaments derived from actin polymers, intermediate filaments and microtubules, which play distinct roles in maintaining cell shape, migration, endocytosis, intracellular transport and the association of cell with the extracellular matrix and cell-cell interactions (Jones et al., 2019). Due to its essential role in these cellular processes, components of the cytoskeleton, particularly the actin cytoskeleton is a common target for infectious agents. For example, *Salmonella enterica* Typhimurium utilizes a set of type III effectors, including SipC, SopE and SptP to reversibly regulate the rearrangement of host actin cytoskeleton to facilitate its entry into non-phagocytic cells (Kubori and Galan, 2003). Other bacterial pathogens such as *Chlamydia*, *Orientia tsutsugamushi*, and *Listeria* also exploit the actin cytoskeleton and microtubule networks to promote their movement in the cytoplasm of host cells and cell to cell spread (Cheng et al., 2018; Grieshaber et al., 2003; Kim et al., 2001).

Growing evidence indicates that manipulation of the actin cytoskeleton dynamics plays an important role in the intracellular lifecycle of *L. pneumophila*. It has been documented that chemical interference of the actin cytoskeleton structure impedes bacterial entry and replication (Charpentier et al., 2009). A number of Dot/Icm effectors have been shown to impose complex modulation of the host actin cytoskeleton. Among these, VipA promotes actin polymerization by functioning as a nucleator (Franco et al., 2012). LegK2 appears to inhibit actin nucleation by phosphorylating the Arp2/3 complex (Michard et al., 2015). The protein phosphatase WipA participate in this regulation by dephosphorylating several proteins involved in actin polymerization, including N-WASP, NCK1, ARP3, and ACK1, leading to dysregulation of actin polymerization (He et al., 2019). RavK is a metalloprotease that cleaves actin in host cells, abolishing its ability to form polymers (Liu et al., 2017). Ceg14 also appears to inhibit actin polymerization by a yet unrecognized mechanism (Guo et al., 2014). Interestingly, LegG1 has been demonstrated to promote microtubule polymerization and host cell migration by functioning as a guanine nucleotide exchange factor (GEF) for the Ran GTPase (Rothmeier et al., 2013; Simon et al., 2014). Counterintuitive to the role of LegG1, cells infected by *L. pneumophila* display defects in migration in a way that requires a functional Dot/Icm system (Simon *et al*., 2014), suggesting the existence of effectors that function to block cell migration.

Herein, we demonstrate that the *L. pneumophila* effector Lem8 (Lpg1290) (Burstein et al., 2009) is a cysteine protease that functions to inhibit host cell migration by targeting the microtubule-associated protein Phldb2 via a mechanism that requires the regulatory protein 14-3-3ζ.

## Results

### Lem8 is a *Legionella* effector with putative cysteine protease activities

One major challenge in the study of bacterial effectors is their unique primary sequences that share little similarity with proteins of known function. Bioinformatics analysis has been proven useful in the identification of putative cryptic functional motifs embedded in their structures. We used PSI-BLAST to analyze a library of the *L. pneumophila* Dot/Icm effectors (Zhu et al., 2011) and found that Lem8 harbors a putative Cys_280_-His_391_-Asp_412_ catalytic triad present in a variety of cysteine proteases (**Fig. 1A**). Further analysis by HHpred (Soding et al., 2005) revealed that Lem8 has high probability to have structural similarity with HopN1 and AvrPphB from *Pseudomonas syringae* (Rodriguez-Herva et al., 2012; Shao et al., 2002), as well as YopT from *Yersinia enterocolitica* (Shao *et al*., 2002) and PfhB1 from *Pasteurella multocida* (Shao *et al*., 2002) (**Fig. S1**).

**Fig. 1.**
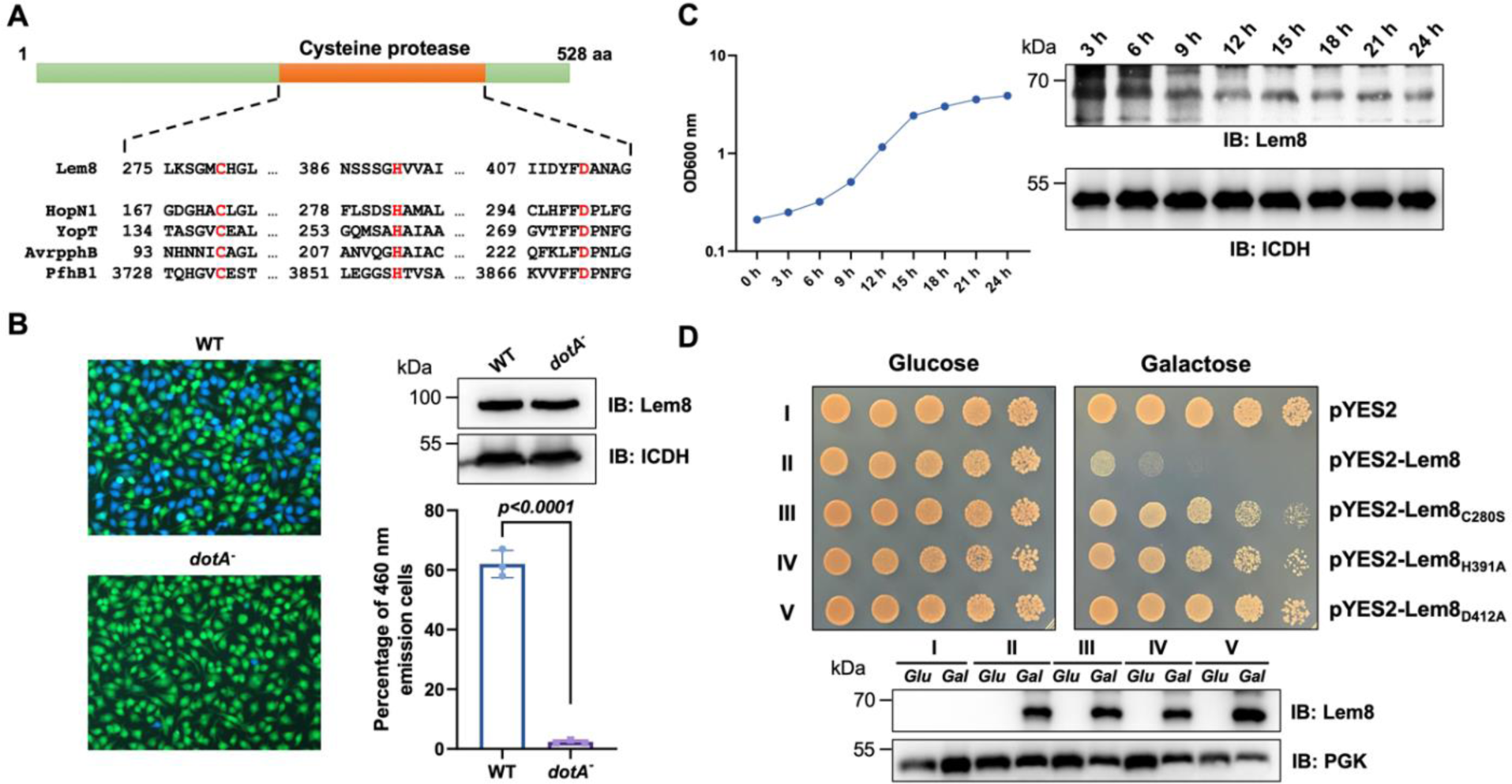
Lem8 is a cysteine protease-like Dot/Icm effector toxic to yeast. **A.** Alignment of Lem8 with several known cysteine proteases obtained by PSI-BLAST analysis. The strictly conserved catalytic residues are marked in red. Shown cysteine proteases are HopN1 and AvrpphB from *P. syringae*, YopT from *Y. enterocolitica*, and PfhB1 from *P. multocida*. **B.** Lem8 is translocated into mammalian cells via the Dot/Icm transporter. U937 cells were infected with wild-type *L. pneumophila* or a *dotA^-^* mutant expressing a β-lactamase- Lem8 fusion. One hour after infection, the CCF4-AM fluorescence substrate was added into the cultures and the cells were incubated for another 2 h at room temperature before image acquisition. Cells emitting blue fluorescence signals were quantitated by counting at least 500 cells in each experiment done in triplicate. Resutls shown are mean ± s.e. from one representative experiment. **C.** Expression profile of *lem8* in *L. pneumophila* grown in AYET broth. Bacteria grown to stationary phase were diluted at 1:20 in fresh medium and subcultures were grown in a shaker. Bacterial growth was monitored by measuring OD_600_ at the indicated time points. Equal amounts of bacterial cells were lysed for measurement of Lem8 levels by immunoblotting with Lem8-specific antibodies. The metabolic protein isocitrate dehydrogenase (ICDH) was probed as loading control. **D.** Lem8 is toxic to yeast in a manner that requires the predicted Cys-His-Asp motif. Yeast strains expressing Lem8 or the indicated mutants from the galactose-inducible promotor were serially diluted and spotted on the indicated media. The plates were incubated at 30°C for 48 h before image acquisition. The expression of Lem8 and its mutants induced by galactose were determined by immunoblotting with Lem8-specific antibodies. The 3- phosphoglycerate kinase (PGK) was detected as loading control.

Lem8 is a protein of 528 residues coded for by the gene lpg1290 in *L. pneumophila* strain Philadelphia 1, it was first identified as a substrate of the Dot/Icm transporter by a machine learning approach (Burstein *et al*., 2009). The translocation of Lem8 by the Dot/Icm system into host cells during *L. pneumophila* infection was later validated by two independent reporter systems (Huang et al., 2011; Zhu *et al*., 2011). Consistent with these results, we observed Dot/Icm-dependent translocation of Lem8 into host cells using the β-lactamase- and CCF2-based reporter assay. Approximately 60% of the cells infected with a Dot/Icm competent strain expressing the β-lactamase-fusion emited blue fluorescence signals. No translocation was detected when the same fusion was expressed in the *dotA*^-^ mutant defective in the Dot/Icm system (Berger and Isberg, 1993) (**Fig. 1B**).

The expression of many Dot/Icm substrates peaks in the post-exponential phase, probably due to the demand for high quantity of effectors to thwart host defense in the initial phase of LCV construction (Luo and Isberg, 2004; Segal, 2013). Thus, we evaluated the expression pattern of *lem8* throughout the entire growth cycle of *L. pneumophila* in broth. Intriguingly, unlike most of effectors, the expression of *lem8* is detected at high levels in the lag phase of its growth cycle in bacteriological medium. A decrease in protein abundance is detected 9 h after the subcultures have started and is maintained constant throughout the remaining 15 h experimental duration (**Fig. 1C**). These results suggest that Lem8 may play a role in the entire intracellular lifecycle of of

### L. pneumophila

Next, we attempted to determine whether the putative cysteine protease motif is important for the effects of Lem8 on eukaryotic cells. We first tested whether Lem8 is toxic to yeast and if so, whether the Cys_280_-His_391_-Asp_412_ motif is required for such toxicity. Expression of Lem8 from the galactose-inducibe promoter caused cell growth arrest (**Fig. 1D**). Mutations in Cys_280_, His_391_ or Asp_412_ did not affect the stability of the protein in yeast, but abolished such toxicity (**Fig. 1D**). Thus, the putative cysteine protease activity conferred by the predicted Cys_280_-His_391_-Asp_412_ catalytic triad very likely is important for the effects of Lem8 on eukaryotic cells.

Genomic analysis reveals that in addition to *L. pneumophila*, *lem8* or its homolog is present only in *L. waltersii*, one of the 40 Legionella species whose genomed had been fully sequenced (Burstein et al., 2016). Such a low prevalence suggests that Lem8 plays a role in the survival of the bacteria in specific inhabits, or its role in other Legionella species is substituted by genes of little sequence similarity that may have arisen by convergent evolution. We probed the role of *lem8* in *L. pneumophila* virulence by examining intracellular replication of the Δ*lem8* mutant in the protozoan host *Dictyostelium discoideum* and in BMDMs. In both host cells, intracellular growth of the Δ*lem8* mutant was indistinguishable to that of the wild-type strain (**Fig. S2**), which is akin to most mutants lacking one single Dot/Icm substrate gene (Qiu and Luo, 2017).

### Lem8 directly interacts with the regulatory protein 14-3-3ζ

To identify the host target of Lem8, we performed a yeast two-hybrid screening against a mouse cDNA Library (Clontech) using Lem8_C280S_ fused to the DNA binding domain of the transcriptional factor GAL4 as bait. Plasmid DNA of the library was introduced into the yeast strain PJ69-4A (James et al., 1996) expressing the bait fusion and colonies appeared on selective medium were isolated and the inserts of the rescued plasmids capable of conferring the interactions were sequenced. We found that 50 out of the 96 independent clones analyzed harbored portions of the gene coding for 14-3-3ζ, a member of a chaperone family important for the activity of a wide variety of proteins in eukaryotic cells (Pennington et al., 2018). Robust interactions occurred in the yeast two-hybrid system when full-length 14-3-3ζ was fused to the AD domain of Gal4 (**Fig. 2A**).

**Fig. 2.**
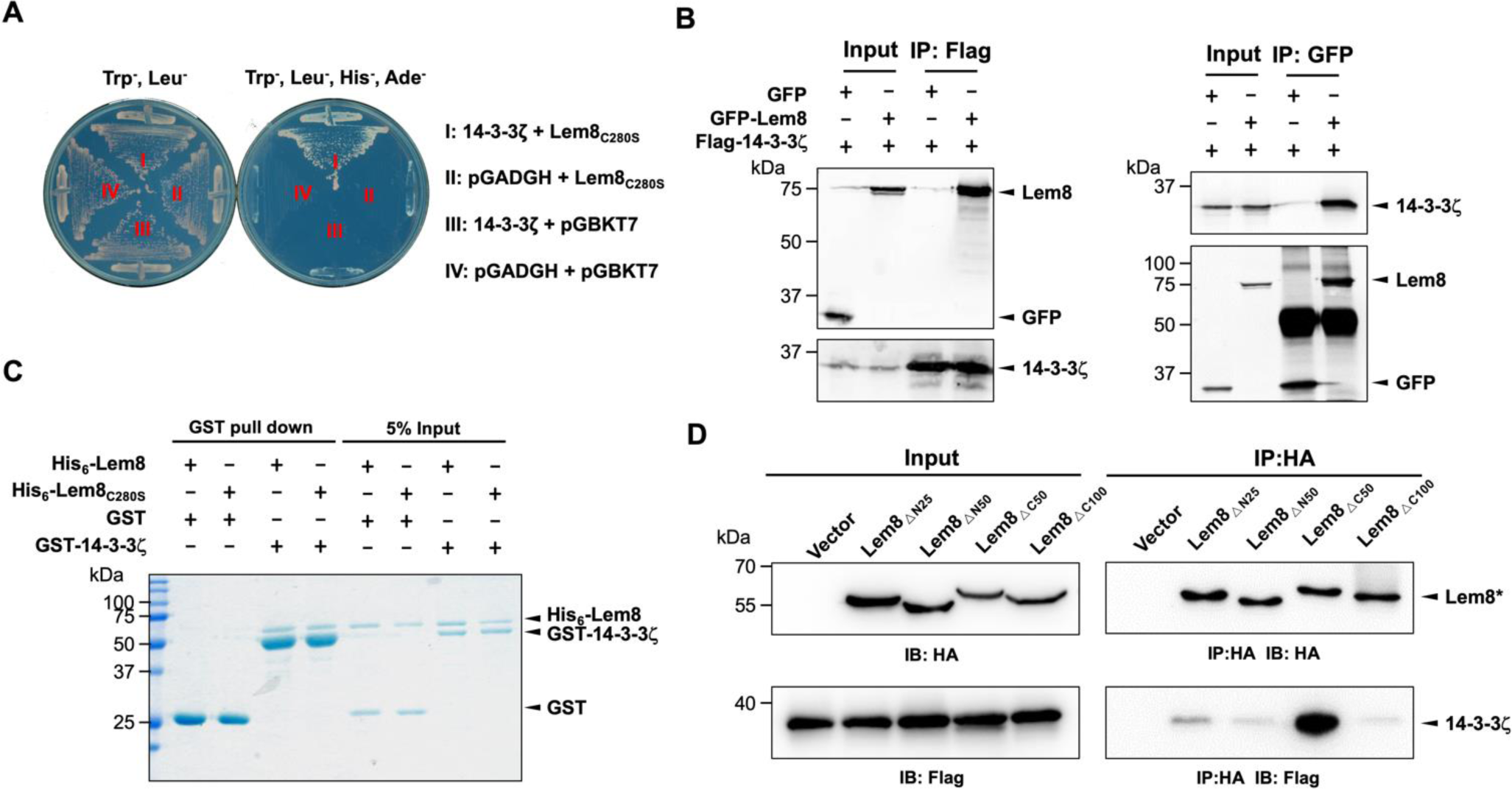
The interactions between Lem8 and 14-3-3ζ. **A.** Interactions between Lem8 and 14-3-3ζ detected by yeast two-hybrid assay. Yeast strains harboring the indicated constructs were streaked on Leu- and Trp- medium to select for plasmids (left) or on Leu^-^, Trp^-^, Ade^-^, and His^-^ medium to assess the interactions (right). Images were acquired after 3-d incubation at 30°C. **B.** Lem8 and 14-3-3ζ form a protein complex in mammalian cell. Total lysates of HEK293T cells transfected with indicated plasmid combinations were immunoprecipitated with a Flag-specific antibody (left panels) or GFP-specific antibodies (right panels), and the precipitates were probed with both Flag and GFP antibodies. Similar results were obtained from at least three independent experiments and the data shown here were from one representative experiment. **C.** Lem8 directly interacts with 14-3-3ζ. GST-14-3-3ζ was incubated with His_6_-Lem8 or His_6_-Lem8_C280S_, and the potential protein complex was captured by glutathione beads for 1 h at 4°C. After extensive washing, bound proteins were solublized with SDS loading buffer, and proteins were detected by Coomassie brilliant blue staining after being resolved by SDS/PAGE. **D.** Interactions between 14-3-3ζ and Lem8 deletion mutants. Lysates of 293T cells expressing Flag-14-3-3ζ and each of the HA-tagged deletion Lem8 were subjected to immunoprecipitation with the anti-HA antibody and the presence of 14-3-3ζ in the precipitates was probed with the Flag-specific antibody.

We further explored the interactions between 14-3-3ζ and Lem8 by reciprocal immunoprecipitation (IP) assays. Flag-tagged 14-3-3ζ was coexpressed with GFP-tagged Lem8 or GFP in HEK293 cells. IP using the Flag antibody specifically precipitated GFP-Lem8, whereas GFP was not detectable in similar experiments. Reciprocally, IP with GFP antibodies specifically pulled down Flag-tagged 14-3-3ζ (**Fig. 2B**). These results suggest that Lem8 forms a complex with 14-3-3ζ in mammalian cells.

To determine whether Lem8 directly binds to14-3-3ζ, we purified recombinant proteins and used GST pulldown assasy to analyze their interactions. We found that mixing His6-Lem8 and GST-14-3-3ζ in reactions led to the formation of stable protein complexes that can be retained by GST beads (**Fig. 2C**).

Members of the 14-3-3 family commonly recognize phospho-serine and/or phospho- threonine sites of client proteins for binding (Muslin et al., 1996). Yet, we did not detect phosphorylation on Lem8 purified from mammalian cells or *E coli* using a pan phospho- serine/threonine antibody. As a control, this antibody detected phosphorylation on CTNNB1, a known phosphorylated target of 14-3-3ζ (Tian et al., 2004). As expected, no signal was detected for ExoS, a non-phosphorylated 14-3-3 interacting effector from *P. aeruginosa* (Henriksson et al., 2002) (**Fig. S3**).

To determine the region in Lem8 involved in binding 14-3-3ζ, we constructed a series of Lem8 deletion mutants and examined their interactions with 14-3-3ζ by immunoprecipitation. Whereas removing as few as 25 residues from the amino terminal end of Lem8 abolished its ability to bind 14-3-3ζ, a Lem8 mutant lacking the last 50 residues can still robustly interact with 14-3-3ζ, and deleting an additional 50 residues from this end abolished the binding (**Fig. 2D**). Thus, either 14-3-3ζ recognizes a large region of Lem8 or deletion from either end of this protein caused significant disruptions in its structure and abolished its ability to interact with 14-3-3ζ.

### Lem8 undergoes 14-3-3ζ-dependent auto-cleavage

Since Lem8 harbors the predicted Cys_280_-His_391_-Asp_412_ catalytic triad associated with proteases from diverse bacterial pathogens, we next investigated whether Lem8 cleaves 14-3-3ζ. Incubation of recombinant His_6_-Lem8 with His_6_-14-3-3ζ at room temperature for 2 h did not lead to detectable cleavage of 14-3-3ζ. Unexpectedly, a protein with a molecular weight slightly smaller than that of Lem8 was detected in this reaction (**Fig. 3A**). The production of this smaller protein did not occur in reactions that contained the Lem8_C280S_ mutatant or when the cysteine protease-specific inhibitor E64 was included in the reactions (**Fig. 3A**), suggesting that this band represents a fragment of Lem8 produced by self-cleavage. Intriguingly, the self-cleavage did not occur in samples containing only Lem8, suggesting that the self-cleavage activity of Lem8 requires 14-3- 3ζ as a co-factor.

**Fig. 3.**
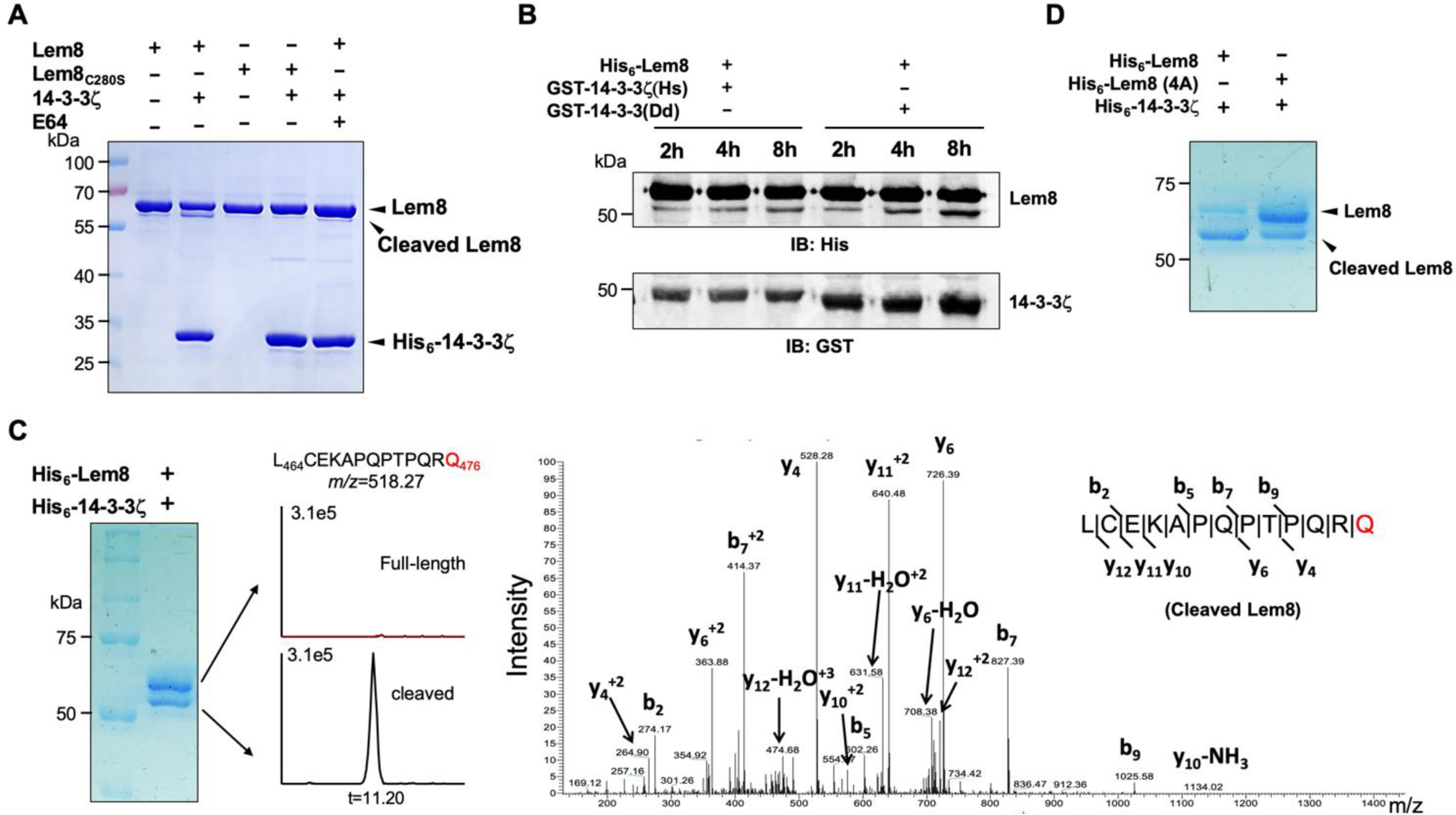
14-3-3ζ induces Lem8 to undergo self-cleavage. **A.** Self-processing of Lem8 requires 14-3-3ζ. His_6_-Lem8 or His_6_-Lem8_C280S_ was incubated with His_6_-14-3-3ζ for 2 h, proteins resolved by SDS-PAGE were detected by Coomassie brilliant blue staining. The cysteine protease inhibitor E64 was added to the indicated samples. **B.** The 14-3-3 protein from *D. discoideum* induces the self-cleavage of Lem8. His_6_-Lem8 was incubated with GST-14-3-3ζ or GST-14-3-3Dd for the indicated time and the mixtures separated by SDS-PAGE were detected by immunoblotting with antibodies specific for Lem8 and GST, respectively. **C.** Determination of the self-cleavage site of Lem8. His_6_-Lem8 was incubated with His_6_- 14-3-3ζ for 16 h, proteins were resolved by SDS-PAGE, stained with Coomassie brilliant blue. Protein bands corresponding to full-length and cleaved Lem8 band was excised, digested with trypsin and analyzed by mass spectrometry. The detection of the semi-tryptic peptide -L_464_CEKAPQPTPQRQ_476_- in cleaved samples suggested that the cleavage site lies between Gln476 and Arg477. **D.** Mutations in cleavage site does not abolish Lem8 self-processing. Recombinant protein of Lem8 and the 4A mutant were each incubated with His_6_-14-3-3ζ for 4 h. Proteins resolved by SDS-PAGE were detected by Coomassie brilliant blue.

*Dictyostelium discoideum*, the protozoan host of *L. pneumophila* codes for one 14- 3-3 protein with 66% identity and 78% similarity to that of human 14-3-3ζ (Eichinger et al., 2005), we investigated whether the *D. discoideum* 14-3-3 (14-3-3Dd) can activate Lem8 by incubating His_6_-Lem8 with GST-14-3-3Dd or human 14-3-3ζ (14-3-3ζHs). In each case, we observed the production of a protein with a size clearly smaller than Lem8 as early as 2 h after the reaction has started. (**Fig. 3B**). Thus, Lem8 can be activated by 14-3-3 from both humans and a protist.

To determine the self-cleavage site of Lem8, we incubated His_6_-Lem8 with His_6_-14- 3-3ζ at room temperature for 16 h. Proteins resolved by SDS-PAGE were stained with Coomassie brilliant blue and bands corresponding to full-length and cleaved Lem8 were excised, digested with trypsin and sequenced by mass spectrometry, repsectively (**Fig. 3C**). Analysis of the tryptic fragments identified a semi-tryptic fragment - A_468_PQPTPQRQ_476_- present in the cleaved protein but not in the full-length protein, suggesting that the cleavage occurs between Gln476 and Arg477 (**Fig. 3C**). To narrow down the potential self-cleavage site, we compared the abundance of identified tryptic peptides from the full-length and cleaved Lem8 and found that the abundance of - A_468_PQPTPQR_475_- was similar between two sets of samples, whereas peptide -A_478_QSLSAETER_487_- was present only in full-length samples but not in the cleaved ones (**Fig. S4A**), suggesting that the cleavage site was between R475 and R487. Consistent with this notion, the signal of a semi-tryptic fragment -L_464_CEKAPQPTPQRQ_476_- was identified in the cleaved protein but not in the full-length protein, suggesting that the cleavage occurs between Gln476 and Arg477 (**Fig. 3C**).

To determine whether Lem8 undergoes auto-cleavage via the recognition of the protein sequence around Gln_476_, we introduced mutations to replace residues Pro_473_, Gln_474_, Arg_475_ and Gln_476_ with alanine and incubated this Lem8 mutant (called 4A) with 14-3-3ζ. Unexpectedly, although at a lower rate, self-cleavage still occurred in this mutant (**Fig. 3D**). We further examined the self-cleavage of Lem8 by fusing GFP to the carboxyl end of Lem8, Lem8_C280S_ and Lem8 _△ C50,_ respectively. These fusion proteins were expressed in HEK293T cells and the cleavage was probed by immunoblotting with GFP-specific antibodies. We found that a fraction of Lem8-GFP and Lem8_△C52_-GFP has lost the GFP portion of the fusions, an event that did not occur in Lem8_C280S_-GFP (**Fig. S4B**). Thus, although the amino acids adjacent to Gln476 play a role in its self-cleavage, other factors such as the overall structure of Lem8 may contribute to the recognition of the cleaving site.

### Lem8 targets Phldb2 for cleavage

It has been reported that some bacterial cysteine proteases cleave both themselves and their substrates in the host by recognizing sites with similar sequences. For instance, AvrpphB and Avrrpt2, two type III effectors from *P. syringae* cleave themselves as well as their host targets PBS1 and RIN4, respectively (Chisholm et al., 2005; Shao et al., 2003). Importantly, in each case, the sequences of the recognition sites for both self-cleavage and cellular target cleavage are very similar. In fact, this feature has been exploited to predict the potential host substrates of these effectors by bioinformatic analyses. Therefore, we performed BLAST searches and obtained 10 candidate proteins that contain sequence elements resembling the self-cleavage site of Lem8, including Phldb2, Rasgrp2, Pak6, Exoc8, Ankrd13B, Chkb, Ppp6R1, Kiaa1033, Gnal and Gpr61.

The predicted recognition sites in these proteins locate in the middle or at sites close to either their amino or carboxyl ends (**Fig. 4A**). Further experiments revealed that one of the candidates, Pleckstrin homology-like domain family B member 2 (Phldb2) can be cleaved by Lem8 in a process that requires an intact Cys_280_-His_391_-Asp_412_ catalytic triad. The predicted Lem8 recognition site locates between the 1106^th^ residue and the 1119^th^ residue in this protein of 1253 amino acids (**Fig. 4A**). In HEK293T cells, expression of Lem8 led to a considerable reduction of endogenous Phldb2 (**Fig. 4B**). To confirm this finding, we added an HA and a Flag tag to the amino and carboxyl end of Phldb2 respectively, and co-expressed the double tagged protein in HEK293T cells with Lem8 or each of the mutants with mutations in one of the three sites (C280S, H391A and D412A) predicted to be critical for catalysis. Detection of tagged Phldb2 by immunoblotting with the Flag-specific antibody indicated that the protein levels in cells expressing Lem8 were reduced comparing to samples in which the catalytically inactive mutants were expressed. Athough to a lesser extent, reduction in Phldb2 was also observed in experiments in which the tagged protein was detected with the HA antibody (**Fig. 4C**).

**Fig. 4.**
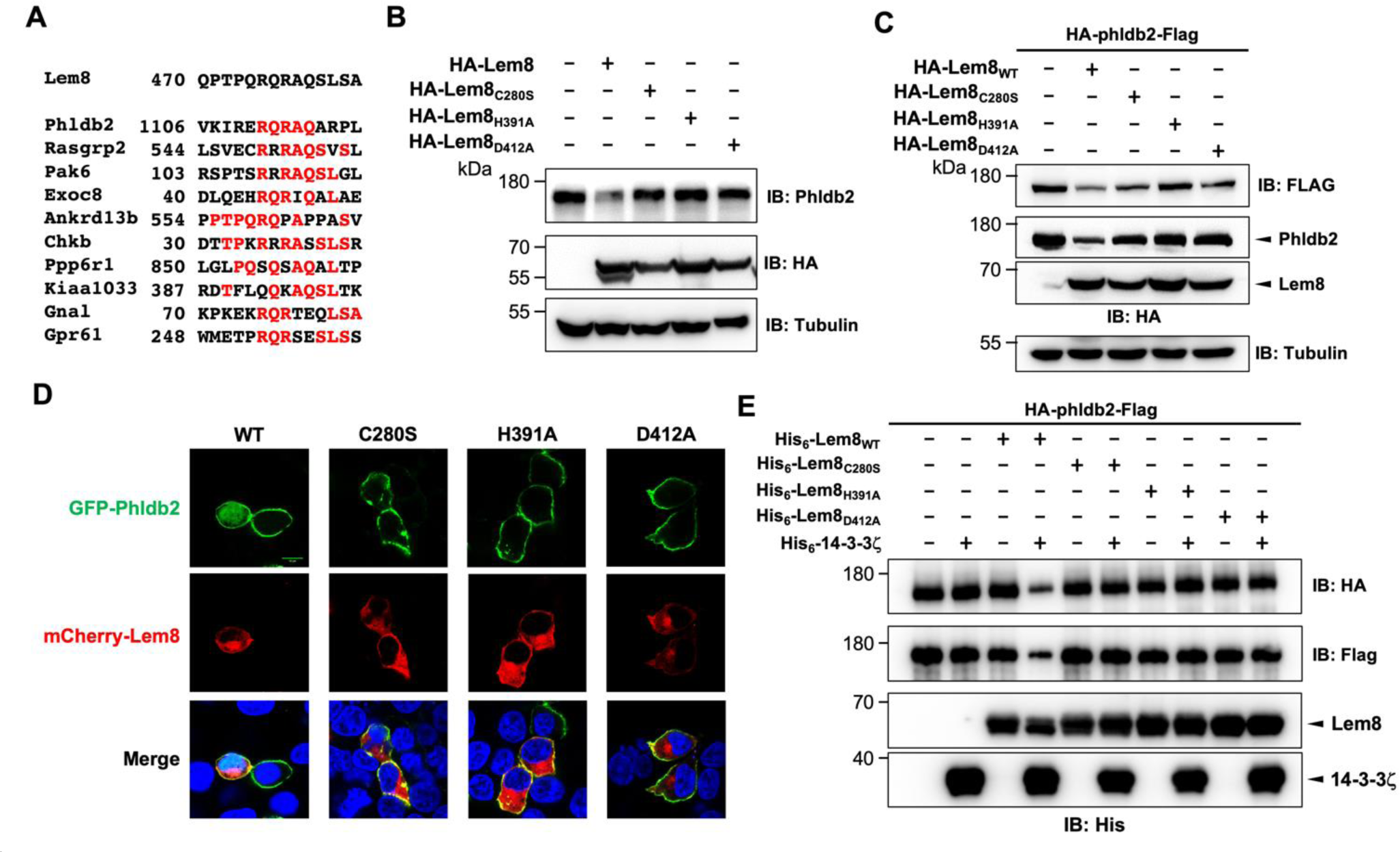
Lem8 cleaves Phldb2 in a manner that requires 14-3-3ζ. **A.** Multiple alignments of the self-cleavage site of Lem8 with potential targets in human cells identified by bioinformatic analysis. Identical residues are highlighted in red. **B.** Lem8 reduces the protein levels of endogenous Phldb2 in mammalian cells. Lem8 and the indicated mutants were individually expressed in HEK293T cells by transfection. 24 h after transfection, the samples were resolved by SDS-PAGE and detected by immunoblotting with anti-Phldb2 antibodies. Tubulin was used as a loading control. Results shown were one representative from three independent experiments with similar results. **C.** Lem8 cleaves exogenous Phldb2 in mammalian cells. HA and Flag tag were fused to the amino and carboxyl end of Phldb2 respectively and the double tagged protein was co-expressed in HEK293T cells with Lem8 or each of the mutants. 24 h after transfection, the samples were resolved by SDS-PAGE and probed by a HA-specific antibody and a Flag-specific antibody, respectively. Tubulin was detected as a loading control. Results shown were one representative from three independent experiments with similar results. **D.** Lem8 alters the subcellular distribution of GFP fused to Phldb2. GFP was fused to the amino end of Phldb2 and the protein was co-expressed in HEK293T cells with mCherry-Lem8 or each of the mutants. 24 h after transfection, cells were fixed and nucleus were stained by Hoechst 33342. The fluorescence Images of GFP (green), mCherry (red) and Hoechst (blue) were acquired with a Zeiss LSM 880 confocal microscope. Bar, 5 μm. **E.** 14-3-3ζ is required for the cleavage of Phldb2 by Lem8. HA-Phldb2-Flag was expressed in HEK293T cells, immunoprecipitated with a Flag-specific antibody, and eluted with 3×Flag peptides. Purified Phldb2 was incubated with His_6_-Lem8 or each of the mutants in reactions with or without His_6_-14-3-3ζ. Total proteins of all samples were resolved with SDS-PAGE, and probed by immunoblotting with a HA-specific antibody, a Flag-specific antibody and a His-specific antibody.

Phldb2 is a phosphatidylinositol-3,4,5-triphosphate (PIP3) binding protein and is associated with the plasma membrane (Paranavitane et al., 2003), we next examined how Lem8-mediated cleavage impacts its cellular localization. In HEK293T cells, when GFP-Phldb2 was ectopically expressed, the GFP signals mainly were associated with the plasma membrane, and this pattern of distribution remains unchanged in cells co-expressing enzymatically inactive Lem8 mutants (**Fig. 4D**). In constrast, in cells co-expressing wild-type Lem8, the GFP signals redistributed to occupy the entire cytoplasm, including the nuclei of transfected cells, a pattern similar to that of GFP itself (**Fig. 4D**). These observations suggest that GFP tag had been cleaved from the GFP-Phldb2 fusion to assume its typical localization in these cells. We also analyzed how Lem8 impacts the subcellular localization of endogenous Phldb2. In cells expressing mCherry-Lem8_C280S_, Phldb2 is mainly associated with the plasma membrane. In contrast, in cells expressing mCherry-Lem8, the association of Phldb2 with the plasma membrane almost became undetectable (**Fig. S5A).**

We next examined whether the cleavage of Phldb2 by Lem8 occurs in a cell-free reaction. HA-Phldb2-Flag expressed in HEK293T cells isolated by immunoprecipitation was incubated with Lem8 or its inactive mutants with or without 14-3-3ζ. Cleavage of Phldb2 occurred only in reactions containing wild-type Lem8 and 14-3-3ζ (**Fig. 4E**). Taken together, these results establish Phldb2 as a target of Lem8.

Our results using the double tagged Phldb2 suggest that Lem8 likely cleaves Phldb2 not only at the predicted site located in the carboxyl end of the protein, but also targets its amino terminal portion (**Fig. 4C-E**). To test this hypothesis, we constructed two Phldb2 mutants by replacing residues Arg_1111_ and Gln_1112_ (Phldb2_AA1_) or Gln_1112_ and Arg_1113_ (Phldb2_AA2_) within the predicted recognition sequence with alanine. Each of these mutants was co-expressed with Lem8 in HEK293T cells by transfection. Comparing to samples expressing enzymatically inactive Lem8, the amounts of protein detected by the amino terminal Flag epitope and the carboxyl end HA tag both decreased in cells co-expressing wild-type Lem8 (**Fig. S5B**). We validate this notion by making constructs in which GFP was fused to the amino terminal end of Phldb2 and three of its truncation mutants, Phldb2_△N50_, Phldb2_△N100_ and Phldb2_△N200,_ respectively. Each of these fusion proteins was co-expressed with Lem8 or Lem8_C280S_ in HEK293T cells and the protein level of these fusions was probed by immunoblotting with GFP-specific antibodies. In each case, a fraction of the protein has lost the GFP portion of the fusions when co-expressed with Lem8 but not with Lem8_C280S_ (**Fig. S5C**). Intriguingly, the cleavage also occurred in fusion proteins in which the GFP is fused to the carboxyl terminus of Phldb2 or Phldb2_△C153_ (**Fig. S5C**). These results are consistent with the notion that Lem8 targets Phldb2 at multiple sites.

We also attempted to detect Lem8-mediated cleavage of endogenous Phldb2 in cells infected with *L. pneumophila*. Although Lem8 translocated into infected cells by a Dot/Icm-competent strain expressing Lem8 from a multicopy plasmid is readily detetable, we were unable to detect Phldb2 cleavage in these samples (**Fig. S5D**). The most likely reason for the inability to detect Lem8 activity against Phldb2 in infected cells is the low abundance or instability of the cleaved protein or a combination of both.

### 14-3-3ζ binds Lem8 by recognizing a coiled-coil motif in its amino terminal region

Using the online MARCOIL sequence analysis software (Gabler et al., 2020), we identified a putative coiled coil motif located in the amino region of Lem8. Coiled coil is a common structural element in proteins, particularly those of eukaryotic origin; it is formed by 2-7 supercoiled alpha-helices (Liu et al., 2006), and often is involved in protein-protein interactions, thus playing an important role in the formation of protein complexes (Burkhard et al., 2001). To determine the role of this region in the activity of Lem8, we introduced mutations to replace Leu_58_ and Glu_59_, the two sites predicted to be essential for the coild coil structure in Lem8, with glycine (called Lem8_GG_) (**Fig. 5A**). When tested in yeast, these mutations have completely abolished the toxicity of Lem8 without affecting its expression or stability (**Fig. 5B** ). These mutations may affect the cysteine protease acitivity of Lem8, its interaction with the regulatory protein 14-3-3ζ or its ability to recognize substrates.

**Fig. 5.**
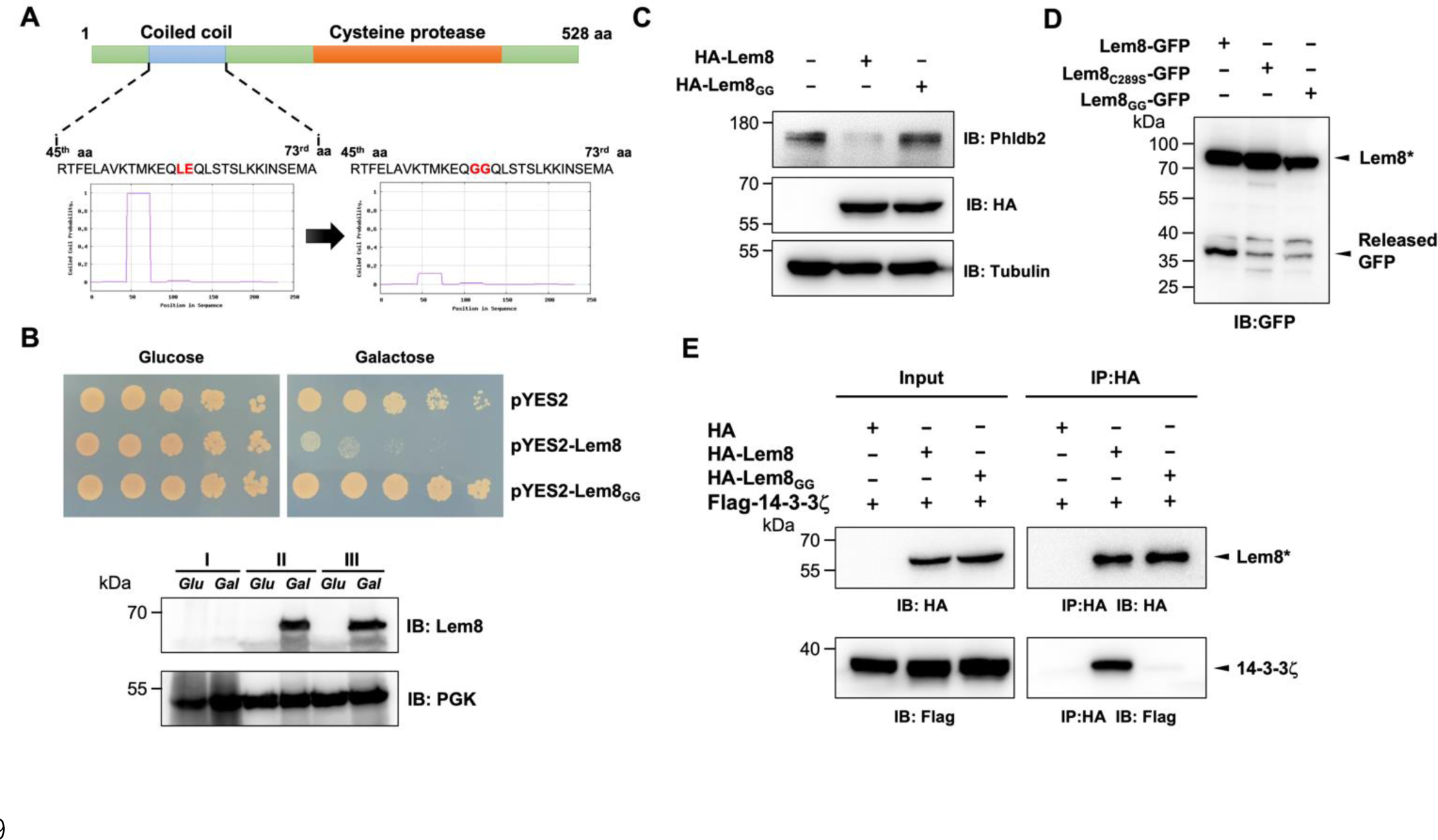
14-3-3ζ binds to Lem8 through a Coiled coil motif. **A.** Lem8 harbors a putative coil motif. A predicted coiled coil motif located in the amino end of Lem8 (top panel). The sequence ranges from the 45^th^ residue to the 73^rd^ residue with a coiled-coil probability of 100% according to MARCOIL (lower panel, left). Replacement of Leu_58_ and Glu_59_ with glycine (highlighted in red) is predicted to reduce the coiled-coil probability to about 10% (lower panel, right). **B.** The predicted coiled coil motif is critical for Lem8-mediated yeast toxicity. Yeast cells inducibly expressing Lem8 or mutant Lem8_GG_ were serially diluted and spotted onto the indicated media for 48 h (top panel). The expression of Lem8 and Lem8_GG_ was examined and PGK1 was probed as a loading control (lower panel). **C.** Lem8_GG_ loses the capacity to cleave Phldb2 in mammalian cells. Lysates of HEK293T cells expressing Lem8 or Lem8_GG_ were resolved by SDS-PAGE and detected by immunoblotting with antibodies specific for Phldb2 and Lem8, respectively. Tubulin was used as a loading control. Results shown were one representative from three independent experiments with similar results. **D.** The predicted coiled coil motif is required for self-processing of Lem8. The indicated alleles of Lem8-GFP were individually expressed in HEK293T cells by transfection. Samples resolved by SDS-PAGE were detected by immunoblotting with GFP-specific antibodies. Results shown were one representative from three independent experiments with similar results. **E.** Interactions between 14-3-3ζ and the Lem8_GG_ mutant. Lysates of 293T cells expressing Flag-14-3-3ζ with HA-Lem8 or HA-Lem8_GG_ were subjected to immunoprecipitation with the anti-HA antibody and the presence of 14-3-3ζ in the precipitates was probed with the Flag-specific antibody.

We examined the ability of Lem8_GG_ to cleave Phldb2 by coexpressing them in HEK293T cells. Whereas wild-type Lem8 consistently cleaves this substrate, Lem8_GG_ has lost such activity despite a similar expression level (**Fig. 5C**). To test the self-cleavage of Lem8_GG,_ we expressed Lem8-GFP or Lem8_GG_-GFP in HEK293T cells and probed the fusion proteins by immunoblotting with GFP-specific antibodies. Comparing to Lem8-GFP, the protein levels of Lem8_GG_-GFP and in Lem8_C280S_-GFP were similarly higher, indicating that the loss of the GFP portion of the fusion occurred in Lem8-GFP but not in Lem8_GG_- GFP (**Fig. 5D**). Finally, we examined the impact of these mutations on the interaction between Lem8 and 14-3-3ζ. Albeit Lem8_GG_ expressed similarly to the wild-type, it has largely lost the ability to bind 14-3-3ζ in immunoprecipitation assays (**Fig. 5E).** Together with the observation that Lem8 mutants lacking as few as 25 residues from its amino terminal end are unable to bind 14-3-3ζ, these results suggest that the regulatory protein most likely bind Lem8 by recognizing the coiled coil motif located in its amino end region.

### Auto-cleaved Lem8 maintains the cysteine protease activity

It has been well-established that some proteins, particularly enzymes are made as precursors or zymogens that need either auto-processing or cleavage by other enzymes to exhibit their biological functions. One such example is caspases involved in cell death regulation and other important cellular functions. These enzymes are synthesized as zymogens before being activated by proteolytic cleavage in response to stimulation (Shalini et al., 2015). In some cases, auto-processing leads to changes or even loss of their enzymatic activity (Kapust et al., 2001; Zhang et al., 2018). To investigate whether Lem8 that has undergone self-cleavage still possesses the cysteine protease activity, we tested the cleavage of Phldb2 by Lem8_△C52_, its self-processed form. Similar to full-length Lem8, Lem8_△C52_ was able to reduced the protein levels of Phldb2. In contrast, other truncation mutants, including Lem8_△N25_, Lem8_△N50_ and Lem8_△C100_ have lost the capacity to cleave Phldb2 (**Fig. 6A**). In addition, Lem8_△C52_, but not Lem8_△N25_ or Lem8_△C100_, cleaved the GFP tag from from the GFP-Phldb2 fusion and released the GFP signals from the plasma membrane (**Fig. 6B**). Intriguigingly, although their ability to cleave Phldb2 appears similar, under our experimental conditions, the protein level of Lem8_△C52_ is considerably lower than that of Lem8 (**Fig. 6A**), suggesting that the self-processed form has higher activity.

**Fig. 6.**
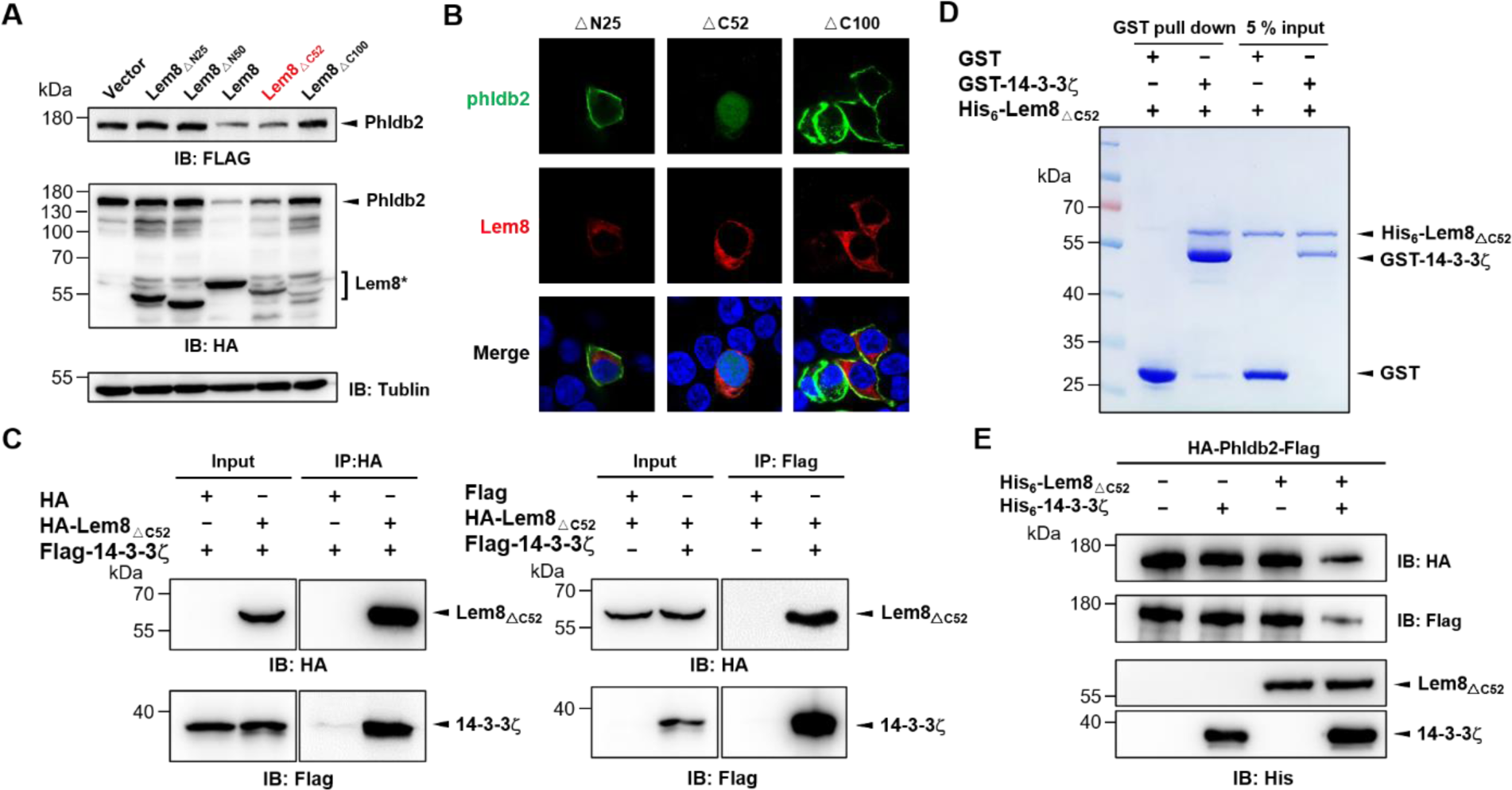
The cysteine protease activity of auto-processing Lem8. **A.** The auto-processed form of Lem8 cleaves Phldb2 in cells. HA-Phldb2-Flag was co- expressed in HEK293T cells with Lem8 or the indicated truncation mutants including the self-processed form, Lem8_△C52_. 24 h after transfection, the samples were resolved by SDS-PAGE and probed by a HA-specific antibody and a Flag-specific antibody. Tubulin was used as a loading control. Results shown were one representative from three independent experiments with similar results. **B.** Lem8_△C52_ causes redistribution of GFP-Phldb2. Truncations of Lem8, including Lem8 _△N25_, Lem8_△C52_ and Lem8_△C100_ fused to mCherry was individually expressed in HEK293T cells with GFP-Phldb2. 24 h after transfection, the fluorescence Images were acquired with a Zeiss LSM 880 confocal microscope. Bar, 5 μm. **C.** The interaction between Lem8 _△ C52_ and 14-3-3ζ. Total lysates of HEK293T cells transfected with indicated plasmid combinations were immunoprecipitated with antibodies specific for HA (left panel) or Flag (right), and the precipitates were probed with both HA and Flag antibodies. Similar results were obtained from at least three independent experiments and the data shown here were from one representative experiment. **D.** Lem8_△C52_ directly interacts with 14-3-3ζ. Mixtures containing GST-14-3-3ζ and His_6_- Lem8_△C52_ were incubated with glutathione beads for 1 h at 4°C. After washing, samples resolved by SDS/PAGE were detected by Coomassie brilliant blue staining. **E.** The cleavage of Phldb2 by Lem8_△C52_ requires 14-3-3ζ. Purified HA-Phldb2-Flag from HEK293T was incubated with His_6_-Lem8_△C52_ in reactions with or without His_6_-14-3-3ζ. Total proteins of all samples were resolved with SDS-PAGE, and probed by immunoblotting with antibody specific for HA, Flag and His_6_, respectively.

We next examined whether the protease activity of Lem8_△C52_ still requires 14-3-3ζ binding. Results from immunoprecipitation and pulldown assays with purified protiens clearly showed that Lem8 _△ C52_ robustly binds 14-3-3ζ (**Fig. 6C-D**). Furthermore, incubation of Lem8_△C52_ with Phldb2 isolated from cells did not lead to its cleavage, but the inclusion of 14-3-3ζ allowed the cleavage to occur (**Fig. 6E**), indicating that Lem8_△C52_ still requires 14-3-3ζ for its protease activity.

### Lem8 inhibits migration in mammalian cells

Phldb2 is a PIP3 binding protein involved in microtubule stabilization (Lansbergen et al., 2006; Paranavitane *et al*., 2003), thus playing a pivotal role in cell motility. Depletion of Phldb2 significantly reduces the migration of MDA-231 cells in the haptotactic migration assay (Astro et al., 2014). As Lem8 cleaves Phldb2, we hypothesized that Lem8 may affect cell migration. To test this, we first established HEK293T-derived cell lines that stably express GFP, GFP-Lem8 or GFP-Lem8_C280S_. Immunoblotting confirmed that Lem8 and Lem8_C280S_ robustly expressed in the respective cell lines. Furthermore, in the cell line expressing Lem8, the level of Phldb2 was drastically reduced comparing to that in the line expressing GFP or Lem8_C280S_ (**Fig. 7A**). We then used the wound-healing scratch assay (De Ieso and Pei, 2018) to examine the impact of ectopic Lem8 expression on cell motility. Confluent monolayer of each cell lines was scratched using a pipette tip and the migration of cells into the gap was monitored over a period of 24 h. Results from this experiment showed that the percentage of wound closure at 24 h after wounding was around 50% in samples using cells expressing GFP or GFP-Lem8_C280S_. In the same experimental duration, cells expressing GFP-Lem8 only filled the gap by 26%, which was significantly slower that of the controls (**Fig. 7B**). Thus, ectopic expression of Lem8 inhibits mammalian cell migration.

**Fig. 7.**
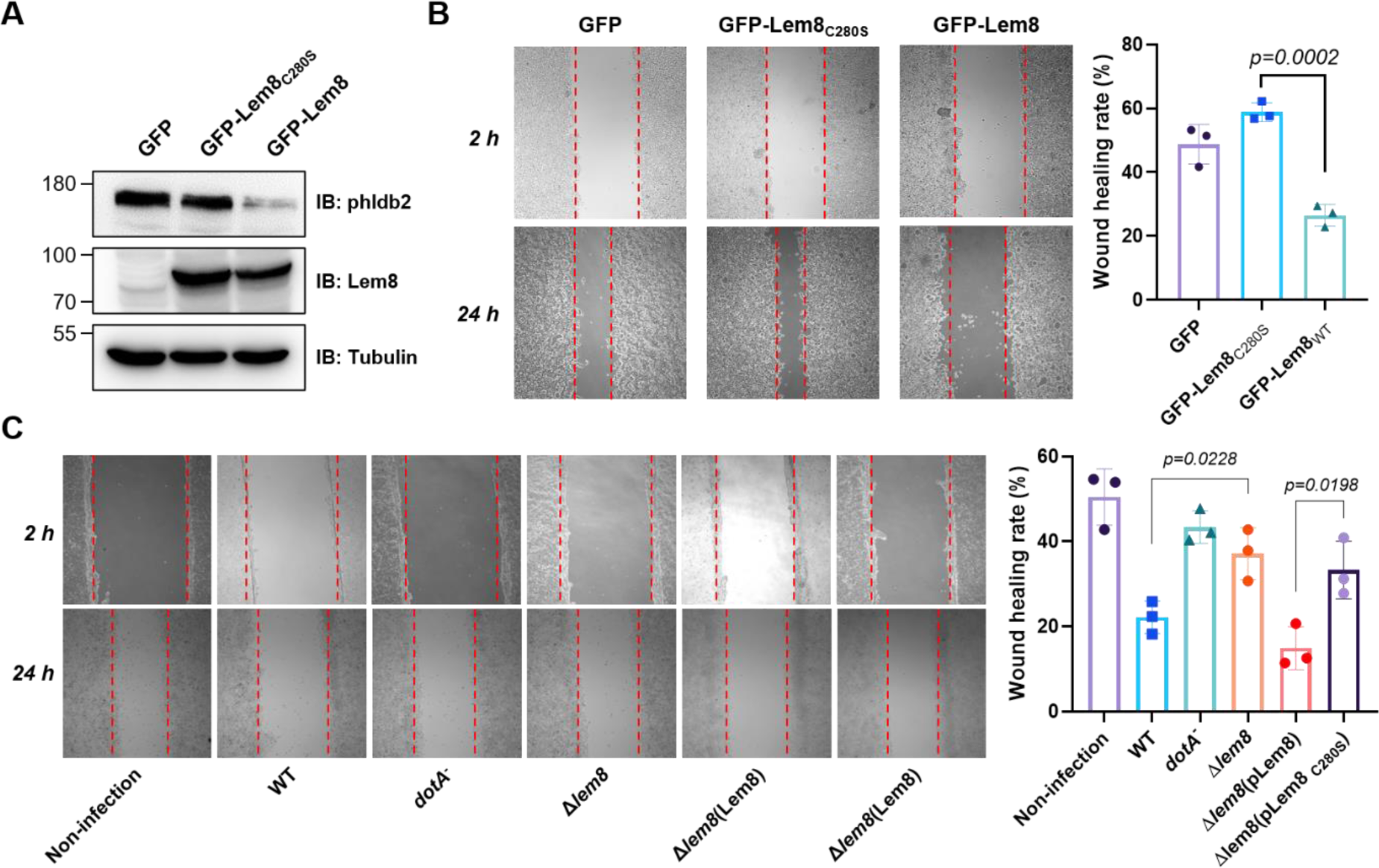
Lem8 contributes to cell migration inhibition by *L. pneumophila*. **A.** Establishment of cell lines stably expressing Lem8 or its enzymatically inactive mutant. HEK293T cells were transduced with lentiviral particles harboring the indicated plasmid at an MOI of 10 for two days, and the GFP-positive cells were isolated by a BD Influx™ cell sorter. Lysates of each cell lines were probed by immunoblotting with antibodies specific for Phldb2 or GFP. Tubulin was used as a loading control. **B.** Wound-healing scratch assay of the three stable cell lines. The three cell lines were individually seeded into 6 well plates. When reached confluency, cell monolayer of each cell lines was scratched using a pipette tip. Images of the wounds were captured using an Olympus IX-83 fluorescence microscope at 2 h and 24 h after making the scratches. Images of a representative experiment were shown (left panel). The wound healing rates from three independent experiments was quantitated by Image J (right panel). **C.** Evaluation of the impact of Lem8 on cell migration in cells infected with *L. pneumophila*. HEK293T cells expressing the FcγII receptor were infected with opsonized bacteria of the indicated *L. pneumophila* strains at an MOI of 50 for 2 h. After washes, the wound- healing scratch assay was performed to evaluate the impact of infection on cell migration. Images of a representative experiment were shown (C, left panel) and the wound healing rate was analyzed by Image J (C, right panel).

An earlier study has shown that in the under-agarose migration assay, *L. pneumophila* inhibits the chemotaxis of mouse macrophages towards cytokines CCL5 and TNF-α in a Dot/Icm-dependent manner (Simon *et al*., 2014). Yet, the Dot/Icm substrates responsible for this inhibition remain elusive. We further studied whether Lem8 contributes to the inhibition of infected cell migration. To test this, we performed the scratch assay with cells infected with wild-type *L. pneumophila* or several strains relevant to *lem8*. The percentage of wound closure by cells infected with wild-type *L. pneumophila* or Δ*lem8*(pLem8) was significantly lower than that with cells infected with the Δ*lem8* mutant. Consistent with its lack of the protease activity, Lem8_C280S_ was unable to complement the defects displayed by the Δ*lem8* mutant (**Fig. 7C**). Thus, the inhibition of cell migration by *L. pneumophila* during infection is caused at least in part by the activity of Lem8.

## Discussion

Intracellular bacteria manipulate cellular processes to create a niche that supports their survival and replication in host cells by virulence factors that target proteins important for the regulation of these processes. These virulence factors often attack host regulatory proteins by diverse posttranslational modifications (PTMs) such as phosphorylation (Krachler et al., 2011), ubiquitination (Zhou and Zhu, 2015), AMPylation (Yarbrough et al., 2009), acetylation (Mukherjee et al., 2006) and ADP-ribosylation (Cohen and Chang, 2018). Proteolytic processing is a type of PTM that can lead to the activation, inactivation or destruction of target proteins, causing alterations in cellular structure or signaling that benefit the pathogen. For instance, the type III effector EspL from enteropathogenic *Escherichia coli* functions as a cysteine protease that antagonizes host inflammatory response by degrading several proteins involved in necroptotic signalling (Pearson et al., 2017). Our results herein establish Lem8 as a cysteine protease that directly targets the microtubule associated protein Phldb2, therefore contributing to the inhibition of host cell migration by *L. pneumophila*. Lem8 joins a growing list of Legionella effectors with protease activity, including the serine protease Lpg1137 that inhibits autophagy by cleaving syntaxin 17 (Arasaki et al., 2017) and the metalloprotease RavK that attacks actin to disrupt the actin cytoskeleton of host cells (Liu *et al*., 2017).

One interesting feature associated with Lem8 is the requirement of 14-3-3ζ for its activity. In line with the notion that amoebae are the primary host of *L. pneumophila*, the sole 14-3-3 protein from *D. discoideum* similarly activates Lem8. In mammals, members of the 14-3-3 family, including 14-3-3ζ often bind their client proteins by recognizing phosphorylated pockets with relatively conserved sequences such as RSX[pS/pT]XP (mode I) and RXXX[pS/pT]XP (mode II) (pS, phospho-serine, pT, phospho-threonine, X, any residue) (Morrison, 2009). Intriguingly, neither of these two motifs is present in Lem8. Consistently, using a pan phospho-serine/threonine antibody capable of detecting phosphorylation of vimentin, another 14-3-3ζ binding protein in mammalian cells, we cannot detect phosphorylation on Lem8 purified from mammalian cells or *E. coli* (**Fig. S3**). The binding of 14-3-3 proteins to unmodified clients is not unprecedented. All isoforms of 14-3-3 bind non-phosphorylated ExoS of *P. aeruginosa* by recongnizing the DALDL element (Henriksson *et al*., 2002), which bears sequence similarity to the unphosphorylated target WLDLE, an artificial R18 peptide inhibitor derived from a phage display library (Petosa et al., 1998). Elements with a sequence similar to these established recogniztion sites are not present in Lem8 nor is there one resembling those in other nonphosphorylated binding targets of 14-3-3, including GPIb-α (Gu and Du, 1998), p75NTR-associated cell death executor (NADE) (Kimura et al., 2001) and CLIC4 (Suginta et al., 2001).

Two lines of evidence suggest that 14-3-3ζ recognizes a coiled coil motif in the amino terminal portion of Lem8. First, deletion of as few as 25 residues from the amino terminus end of Lem8 abolished its interaction with 14-3-3ζ (**Fig. 2D**). Second, the integrity of a predicted coiled coil motif in the amino terminal portion of Lem8 is required for its binding to the regulatory protein (**Fig. 5**). Coiled coil motifs have long been known to be important for protein-protein interaction but its involvement in binding 14-3-3 has not yet been established. The binding of 14-3-3 to TRIM25 had been suggested to be mediated by recognizing a coiled coil domain, but the mechanism of such binding or whether phosphorylation is required remains unclear (Gupta et al., 2019). Future study, particularly structural analysis of the Lem8-14-3-3ζ complex may allow a definite identification of the region in Lem8 recognized by 14-3-3ζ, which will surely shed light on the additional features of the sequences recognizable by these imporant regulatory proteins.

The self-cleavage of Lem8 has allowed us to identify its recognition sequence and several candidate cellular targets. One unexpected observation is that mutations in the identified recogniztion element reduced but did not abolished self-cleavage (**Fig. 3D**). Thus, the primary sequence may not the only factor that dictates the specificity of Lem8 in substrate recognition. Other factors such as the overall structure of substrates may contribute to the determination of the cleavage site. The low level of promiscuity in cleavage site selection may allow Lem8 to more effectively to bring down the protein level of its cellular targets, which may explain the requirement of 14-3-3ζ for its activity. If a host co-factor is not needed for its activity, Lem8 may cleave itself or even other proteins in *L. pneumophila* cells. For Lem8, self-cleavage in the absence of 14-3-3ζ in bacterial cells will be disastrous because the cleaved product will lose the portion of the protein that harbors translocation signals recognized by the Dot/Icm system (Luo and Isberg, 2004; Nagai et al., 2005). Likewise, the requirement of CaM by the Dot/Icm effector SidJ to inhibit the activity of members of the SidE family is to ensure that such inhibition does not occur in bacterial cells (Bhogaraju et al., 2019; Black et al., 2019; Gan et al., 2019; Sulpizio et al., 2019). The promiscuity in cleavage site recognition by Lem8 is also supported by the observation that this protease appears to cleave Phldb2 at multiple sites (Figs. 4 and S5).

Interference with host cell motility appears to be a common strategy used by bacterial pathogens. For example, *Salmonella enterica* Typhimurium inhibits the migration of infected macrophages and dendritic cells in a process that requires its type III effector SseI, which binds to IQGAP1, an important regulator of cell migration (McLaughlin et al., 2009). Similarly, the phosphatidylinositol phosphatase IpgD from *Shigella flexneri* contributes to the inhibition of chemokine-induced migration of human T cells (Konradt et al., 2011). The observation that cells infected with the wild-type *L. pneumophila* or strain Δ*lem8*(pLem8) migrated significantly slower than those infected with the Δ*lem8* mutant or its complementation strain expressing the Lem8_C280A_ mutant suggests a role of Lem8 in cell mobility inhibition (Simon *et al*., 2014).

Akin to most *L. pneumophila* Dot/Icm effectors, Lem8 is not required for proficient bacterial intracellular growth in commonly used laboratory hosts such as *D. discoideum*. Lem8 may be required for the survival of the bacteria in some specific inhabits or other Dot/Icm effectors may substitute its role by distinct mechanisms, thus contributing to such inhibition. Future studies aiming at the identification and characterization of Dot/Icm effectors involved in attacking host cells motility will continue to provide insights into the mechanisms of not only bacterial virulence but also the regulation of eukaryotic cell migration.

## Materials and Methods

### Bacterial stains, plasmids and cell culture

*E. coli* strain DH5α was used for plasmid construction and strain BL21(DE3) or XL1blue was used for recombinant protein production and purification. All *E. coli* strains were grown on LB agar plates or in LB broth at 37°C. For maintenance of plasmids in *E. coli*, antibiotics were added in media at the following concentrations: ampicillin,100 μg/mL; kanamycin, 30 μg/mL. All *L. pneumophila* strains were derived from the Philadelphia 1 strain Lp02 and the *dotA^-^* mutant strain Lp03 (Berger and Isberg, 1993) and were listed in **Table S1**. *L. pneumophila* was cultured in N-(2-acetamido)-2-aminoethanesulfonic acid buffered yeast extract medium (AYE) or on charcoal buffered yeast extract plates (CYE). When necessary, thymidine was added into AYE at a final concentration of 0.2 g/mL. pZL507 and its derivatives which allow expression of His_6_-tagged proteins (Xu et al., 2010) in *L. pneumophila* were maintained by thymidine autotrophic. Deletion of the Lem8 coding gene lpg1290 from the genome of *L. pneumophila* was performed as described previously (Liu and Luo, 2007).

Plasmids used in this study are listed in **Table S1**. Genes were amplified by polymerase chain reactions (PCR) using Platinum™ SuperFi II Green PCR mix (Invitrogen, cat# 12369050). The PCR product was digested with restriction enzymes (New England Biolabs, NEB), followed by ligated to linearized plasmid using T4 DNA ligase (NEB). For site-directed mutagenesis, plasmid was reacted with primer pairs designed to introduce the desired mutations using Quikchange kit (Agilent, cat# 600670). After digestion with the restriction enzyme *Dpn*I (NEB, cat# R0176), the products were transformed into *E. coli* strain DH5α. All substitution mutants were verified by double strand DNA sequencing. The sequences of primers used for molecular cloning are listed in **Table S1**.

HEK293T and Hela cells purchased from ATCC were cultured in Dulbecco’s modified minimal Eagle’s medium (DMEM) supplemented with 10% Fetal Bovine Serum (FBS). Bone marrow cells were isolated from 6- to 10-week-old female A/J mice (GemPharmatech, Co., Ltd.) and were differentiated into BMDMs using L929-cell conditioned medium as described previously (Conover et al., 2003). PCR-based test (Sigma, cat# MP0025) was used to validate the absence of potential mycoplasma contamination in all mammalian cell lines. pAPH-HA, a derivative of pVR1012 (Wang et al., 2018) suitable for expressing proteins with an amino HA tag and a carboxyl Flag tag.

### Yeast manipulation

Unless otherwise indicated, yeast strains used in this study were derived from W303 (Thomas and Rothstein, 1989); yeast was grown at 30°C in yeast extract, peptone, dextrose medium (YPD) medium or in appropriate amino acid dropout synthetic media supplemented with 2% of glucose or galactose as the sole carbon source.

For assessment of inducible protein toxicity, Lem8 or its derivatives were cloned into pYES2/NTA (Invitrogen) in which their expression is driven by the galactose-inducible promoter. Yeast transformation was performed using the lithium acetate method (Gietz et al., 1995). After growing in selective liquid medium with 2% raffinose, yeast cultures were serially diluted (five-fold) and 10 μL of each dilution was spotted onto selective plates containing glucose or galactose. Plates were incubated at 30°C for 3 days before image acquisition.

To screen Lem8-interacting protein(s), Gal4-based two-hybrid screening against the mouse cDNA library (Clontech) was performed as described before (Mitsuzawa et al., 2005). Briefly, Lem8_C280S_ was inserted into pGBKT7 (Banga et al., 2007) to give pGBKLem8, which was transformed into the yeast strain PJ-64A (James *et al*., 1996) and the resulting strain was used for yeast two-hybrid screening. The mouse cDNA library was amplified in accordance with the manufacturer’s instructions and the plasmid DNA was transformed into strain PJ-64A (pGBKLem8). Transformants were plated onto a selective synthetic medium lacking adenine, tryptophan, leucine, and histidine, ccolonies appeared on the selective medium were verified for interactions by reintroducing into strain PJ-64A (pGBKLem8) and inserts of those that maintained the interaction phenotype were sequenced to identify the interacting proteins.

To validate the interactions between 14-3-3ζ and Lem8, its full-length gene was inserted into pGADGH (Banga *et al*., 2007) and the plasmids were introduced yeast strain PJ-64A (pGBKLem8). Yeast strains harboring the indicated plasmid combinations were streaked on Leu_-_ and Trp^-^ synthetic medium to select for plasmids and the transformants were transferred to Leu^-^, Trp-, Ade^-^, and His^-^ medium to examine protein-protein interactions measured by cell growth.

### Antibodies and immunoblotting

Polyclonal antibody against Lem8 were generated according to the protocol described before (Guide for the Care and Use of Laboratory Animals, 1996; J. Derrell Clark, 1997). Briefly, 1 mg of emulsified His6-Lem8 with complete Freund’s adjuvant was injected intracutaneously into a rabbit 4 times at 10-day intervals. Sera of the immunized rabbit containing Lem8-specific antibodies were used for affinity purification of IgG with an established protocol (Harlow, 1999).

Samples from cells or bacteria lysates were prepared by adding 5×SDS loading buffer and heated at 95°C for 10 min. The soluble fraction of the lysates was resolved by SDS-PAGE, proteins were transferred onto polyvinylidene fluoride (PVDF) membranes (Pall Life Sciences). The membranes were blocking with 5% nonfat milk for 30 min, followed by incubated with primary antibodies at the indicated dilutions: α-Phldb2 (Sigma, cat# HPA035147, 1:1000), α-HA (Sigma, cat# H3663, 1:3000), α-Flag (Sigma, Cat# F1804, 1: 3000), α-GFP (Proteintech, cat# 50430-2-AP, 1:5000), α-GST (Proteintech, cat# 66001-2, 1:10000), α-His (Sigma, cat# H1029, 1: 3,000), α-ICDH (1: 10,000) (Xu *et al*., 2010), α-Lem8 (1: 5,000), α-PGK (Abcam, cat# ab113687, 1:2,500) and α-Tubulin (Bioworld, cat# AP0064, 1:10,000). After washed 3 times, the membranes were incubated with appropriate HRP-labeled secondary antibodies and the signals were taken and analyzed by Tanon 5200 Chemiluminescent Imaging System.

### Transfection and immunoprecipitation

When grown to approximately 80% confluence, HEK293T cells were transfected using Lipofectamine 3000 (Invitrogen, cat# L3000150) according to the manufacturer’s protocol. Twenty-four hours after transfection, cells were lysed using a lysis buffer (50 mM Tris-HCl, 150 mM NaCl, 0.5% Triton X-100, PH 7.5) for 10 min on ice, followed by centrifugation at 12,000*g* at 4°C for 10 min. Beads coated with Flag- (Sigma, cat# F2426), HA- (Sigma, cat# E6779) or GFP-specific antibodies (Sigma, cat# G6539) were washed twice with lysis buffer and then mixed with the prepared cell supernatant. The mixture was incubated on a rotatory shaker at 4°C overnight. The resin was washed with the lysis buffer for five times, followed by boiling in the Laemmli buffer at 95°C for 10 min to release the bound Flag- or HA-tagged proteins. For proteins used in biochemical reactions, the Flag- or HA-tagged proteins were eluted with Flag peptide (Sigma, cat# F4799) or HA peptide (Sigma, cat# I2149), respectively.

### Protein expression and purification

Lem8 and its mutants were amplified by PCR and cloned into pQE30 to express His_6_-tagged proteins. The plasmids were transformed into in *E. coli* strain XL1blue and grown in LB broth. When the cell density reached an OD_600_ of 0.8, isopropyl-β-D-thiogalactopyranoside (IPTG) was added into the cultures at a final concentration of 0.2 mM to induce the expression of target proteins for 14 h at 16°C. Cells collected by centrifugation were re-suspended in a lysis buffer (1×PBS, 2 mM DTT and 1 mM PMSF), and were lysed with a cell homogenizer (JN-mini, JNBIO, Guangzhou, China). The lysates were centrifugated at 20,000*g* for 30 min at 4°C twice to remove cell debris. The supernatant was incubated with Ni^2+^-NTA beads (QIAGEN) at 4°C for 1 h, followed by washed with 50x bed volumes of 20 mM imidazole to remove unbound proteins. The His6-tagged proteins were eluted with 250 mM imidazole in PBS buffer. Purified proteins were dialyzed in a storage buffer (30mM NaCl, 20 mM Tris, 10% glycerol, pH 7.5) overnight at 4°C and then stored at -80°C.

14-3-3ζ and its homologous genes were cloned into pGEX6p-1 to express GST-tagged proteins. The plasmids were transformed into *E. coli* strain BL21(DE3). Protein expression induction and purification was carried out similarly with Glutathione Sepharose 4B (GE Healthcare) beads. The resin was collected and washed for with wash buffer (lysis buffer plus 200 mM NaCl). The GST-tagged proteins were eluted with 10 mM glutathione and stored at -80°C after dialysis.

### *In vitro* cleavage assays

For auto-cleavage assays, 5 μg His6-Lem8 or its mutants was incubated with or without 2.5 μg 14-3-3ζ in 50 μl reaction buffer (50 mM Tris, 150 mM NaCl, PH 7.5) at room temperature for the indicated time points. For Phldb2 cleavage, Flag-Phldb2 purified from HEK293T cells were added into reactions with or without Lem8 and 14-3- 3ζ at room temperature for the indicated time. In each case, samples were analyzed by SDS-PAGE followed by immunoblotting or Coomassie brilliant blue staining.

### GST pulldown assay

GST-14-3-3ζ or GST bound to Glutathione Sepharose 4B was incubated with His_6_- Lem8 in a binding buffer (50 mM Tris, 137 mM NaCl, 13.7 mM KCl) for 2 h at 4 °C. After washing three times with the binding buffer, beads were boiled in the Laemmli buffer at 95°C for 10 min and the samples were resolved by SDS-PAGE. Proteins were detected by Coomassie brilliant blue staining.

### Bacterial infection, immunostaining and image analysis

For infection experiments, *L. pneumophila* strains were grown in AYE broth to the post-exponential growth phase (OD_600_=3.3-3.8). When necessary, complementation strains were induced by 0.1 mM IPTG for another 4 h at 37°C before infection.

To determine intracellular bacterial growth, *D. discoideum* or BMDMs of A/J mice were infected with relevant *L. pneumophila* at a multiplicity of infection (MOI) of 0.05. 2 h after adding the bacteria, the cells were washed using warm PBS to remove the extracellular bacteria. *D. discoideum* and BMDMs were maintained in 22°C and 37°C, respectively. At the indicated time points, cells were lysed with 0.2% saponin and appropriately diluted lysates were plated on CYE plates. After 4-day incubation at 37°C, the counts of bacterial colonies were calculated to evaluate the growth.

To determine the impact of the infection on cell migration, HEK293T cells transfected to express FcγRII receptor (Qiu et al., 2016) were infected with the indicated bacterial strains. 2 h after infection, cells were washed using warm PBS and were used for the wound healing assay.

To determine the cellular localization of Lem8 in infected cells, BMDMs were infected with relevant *L. pneumophila* strains at an MOI of 10 for 2 h. The samples were immunostained as described earlier (Haenssler et al., 2015). Briefly, we washed the samples 3 times with PBS to remove extracellular bacteria, and fixed the cells with 4% paraformaldehyde at room temperature for 10 min. After three times washes, cells were permeabilized using 0.1% Triton X-100 and then were blocked with 4% goat serum for 1 h. Samples were incubated with rat anti-*Legionella* antibodies (1:10,000) and rabbit anti- Lem8 antibodies (1:100) at 4°C overnight. followed by incubated with appropriate fluorescence-labeled secondary antibodies at room temperature for 1 h. After stained by Hoechst 33342 (Invitrogen, cat# H3570, 1:5000), samples were inspected using an Olympus IX-83 fluorescence microscope.

To detect the cleavage of endogenous Phldb2 by Lem8, Hela cells transfected to express mCherry-Lem8 or mCherry-Lem8_C280S_ were stained with Phldb2-specific antibodies (1:100) as described above. The images were taken using a Zeiss LSM 880 confocal microscope. The determine the impact on ectopically expressed Phldb2, mCherry-Lem8 or mutants each was co-transfected with GFP-Phldb2 into HEK293T cells seededonto glass coverslips (Nest, cat# 801001). Fixed samples were stained with Hoechst, cell images were acquired by a confocal microscope.

### Production of lentiviral particles and transduction

For production of lentiviral particles carrying *lem8* or its mutants, the *gfp*-*lem8* fusion was inserted into pCDH-CMV-MCS-EF1a-Puro (System Biosciences, cat# CD510B-1). The plasmids were co-transfected with pMD2.G (gift from Dr. Didier Trono, Addgene#12259) and psPAX2 (gift from Dr. Didier Trono, Addgene #12260) into HEK293T cells grown to about 70% confluence. Supernatant was collected after 48 hours incubation, followed by filtration with 0.45-μm syringe filters. After measuring the titers using qPCR with the Lentivirus Titer Kit (abm, cat# LV900), the packed lentiviral particles were used to infect newly prepared HEK293T cells at an MOI of 10. After incubation for 2 days, cells were sorted by BD Influx™ cell sorter to establish cell lines stably expressing the gene of interest.

### Mass spectrometry analysis of Lem8 self-cleavage site

Recombinant His_6_-Lem8 was incubated with His_6_-14-3-3 for 8 h and the samples were separated by SDS-PAGE. After Coomassie brilliant blue staining, bands corresponding full-length His_6_-Lem8 or cleaved were excised and subjected to in-gel digestion with trypsin. Peptides were loaded into a nano-LC system (EASY-nLC 1200, Thermo Scientific) coupled to an LTQ-Orbitrap mass spectrometer (Orbitrap Velos, Thermo Scientific). Peptides were separated in a capillary column (75 µm x 15 cm) packed with C18 resin (Michrom BioResources Inc., 4 µm, 100 Å) with the following gradient: solvent B (100 ACN, 0.1% FA) was started at 7% for 3 min and gradually raised to 35% in 40 min, then rapidly increased to 90% in 2 min and maintained for 10 min before column equilibration with 100% solvent A (97% H_2_O, 3% ACN, 0.1% FA). The flow rate was set at 300 nL/min and eluting peptides were directly analyzed in the mass spectrometer. Full-MS spectra were collected in the range of 350 to 1500 *m/z* and the top 10 most intense parent ions were submitted to fragmentation in a data-dependent mode using collision-induced dissociation (CID) with the max injection time of 10 milliseconds. MS/MS spectra were searched against the *Legionella pneumophila* (strain Philadelphia 1) database downloaded from UniProt using Mascot (Matrix Science Inc.). The signals of Lem8 tryptic peptides were compared between full-length and cleaved samples to narrow down the potential cleavage site(s) within specific peptides, cleaved Lem8 semi-tryptic peptides were inspected manually.

### Wound healing assay

Wound healing assays were performed as previously described (Liang et al., 2007). Briefly, HEK293T cells were seeded into 6-well plates and incubated until the confluency reached about 90%. The cell monolayer was scraped in a straight line using a p200 pipet tip to create a “wound”, followed by washing with growth medium to remove the debris. Reduced-serum medium (1% serum) was added and the cells were placed back in a 37°C incubator. 2 h and 24 h after making the scratch, images of the cell monolayer wound were taken using an Olympus IX-83 fluorescence microscope. For each image,distances between one side of the wound and the other were quantitated by Image J (http://rsb.info.nih.gov/ij/). The wound healing rate was calculated by the following formula: % wound healing = (0 h distance – 24 h distance)/24 h distance × 100.

### Data quantitation, statistical analyses

All data were represented as mean ± standard deviation (SD). Student’s *t*-test was applied to analyze the statistical difference between two groups each with at least three independent samples.

## Acknowledgements

The authors thank Dr. Shaohua Wang for plasmids, the study was funded in part by Jilin Science and Technology Agency grant 20200403117SF (LS), 20200901010SF (DL), National Natural Science Foundation of China grant 21974002 (XL), Beijing Municipal Natural Science Foundation grant 5202012 (XL), and the National Institutes of Health grant R01AI127465 (ZQL).

## Author contributions

LS, ZQL, YL and YT conceived the projects, LS, YL, YT, JL, DH, and YZ performed the experiments. KY and XL performed the mass spectrometric analysis. SL, YT, YL, XL, and ZQL analyzed data. SL drafted the first version of the manuscript, and ZQL revised the manuscript with input from all authors.

---

**Fig. S1.**
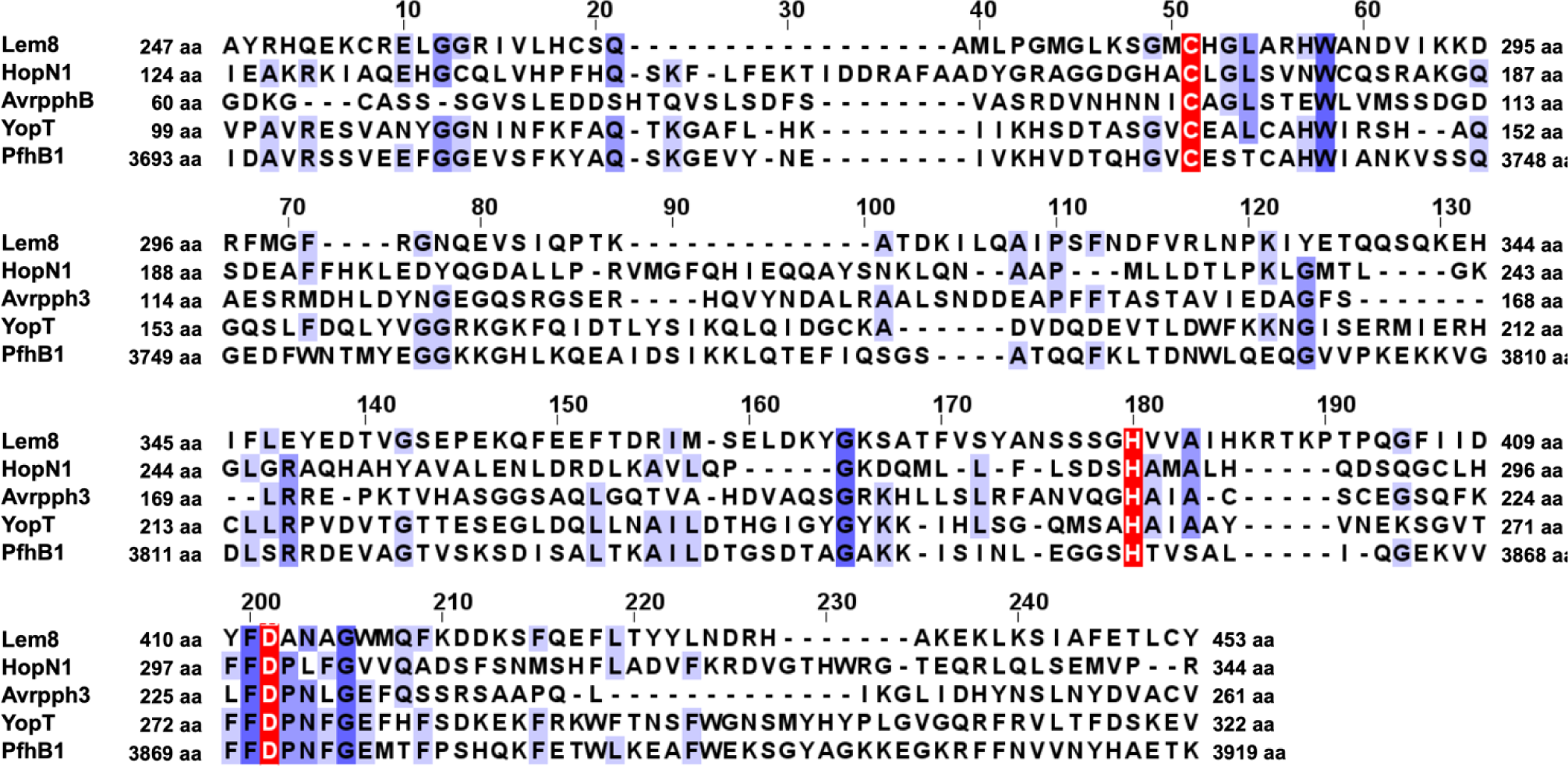
Sequence alignment of Lem8 with four bacterial cysteine protease effectors. The strictly conserved residues were shown in dark purple background and the Cys-His-Asp motif are marked with white letters in a red background. HopN1 and AvrpphB are from *P. syringae*. YopT and PfhB1 are from *Y. pestis* and *P. multocida*, respectively.

**Fig. S2.**
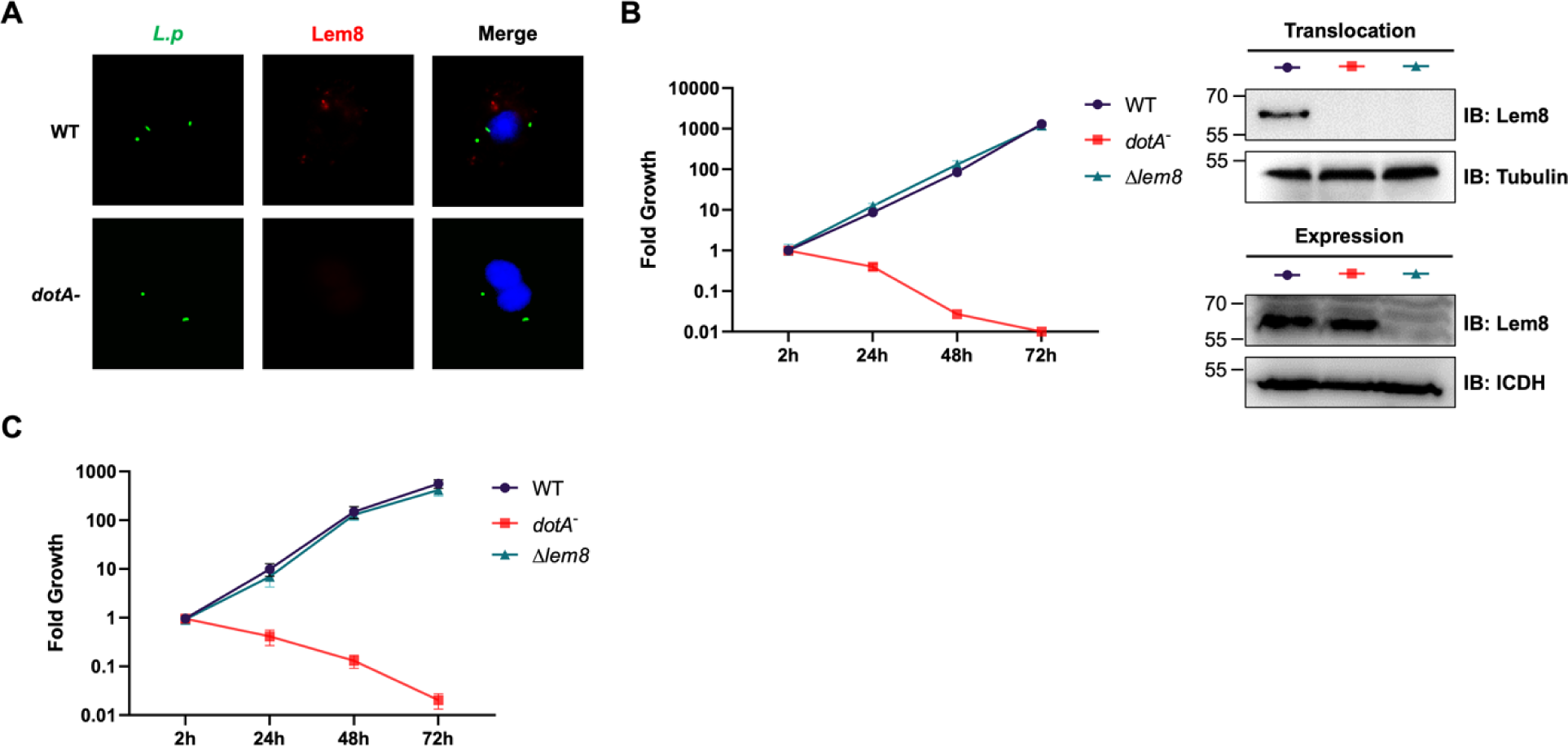
Lem8 is dispensable for intracellular growth of *L. pneumophila*. **A.** Subcellular distribution of Lem8. 2h after infected with the indicated bacterial strains, BMDMs were immunostained using anti-Legionella antibody to identify the bacterial vacuoles (green), followed by staining with the Lem8 specific antibody (red). The nucleus was stained using Hochest 33342 (blue). **B.** Intracellular growth of the Δ*lem8* strain in *D. discoideum*. *D. discoideum* were infected with the indicated bacterial strains at an MOI of 0.1, and the intracellular growth was determined at a 24-h interval for 72 h (left panel). The expression and translocation of Lem8 in each strain was probed with Lem8 specific antibodies (right panel). ICDH and Tubulin were used as loading controls for bacterial and host cells, respectively. Similar results were obtained in three independent experiments. **C.** Intracellular growth of Δ*lem8* strain in BMDMs. The bacterial strains were used to infect BMDMs at an MOI of 0.1 and the intracellular growth was monitored at the indicated time points (left panel). Similar results were obtained in three independent experiments.

**Fig. S3.**
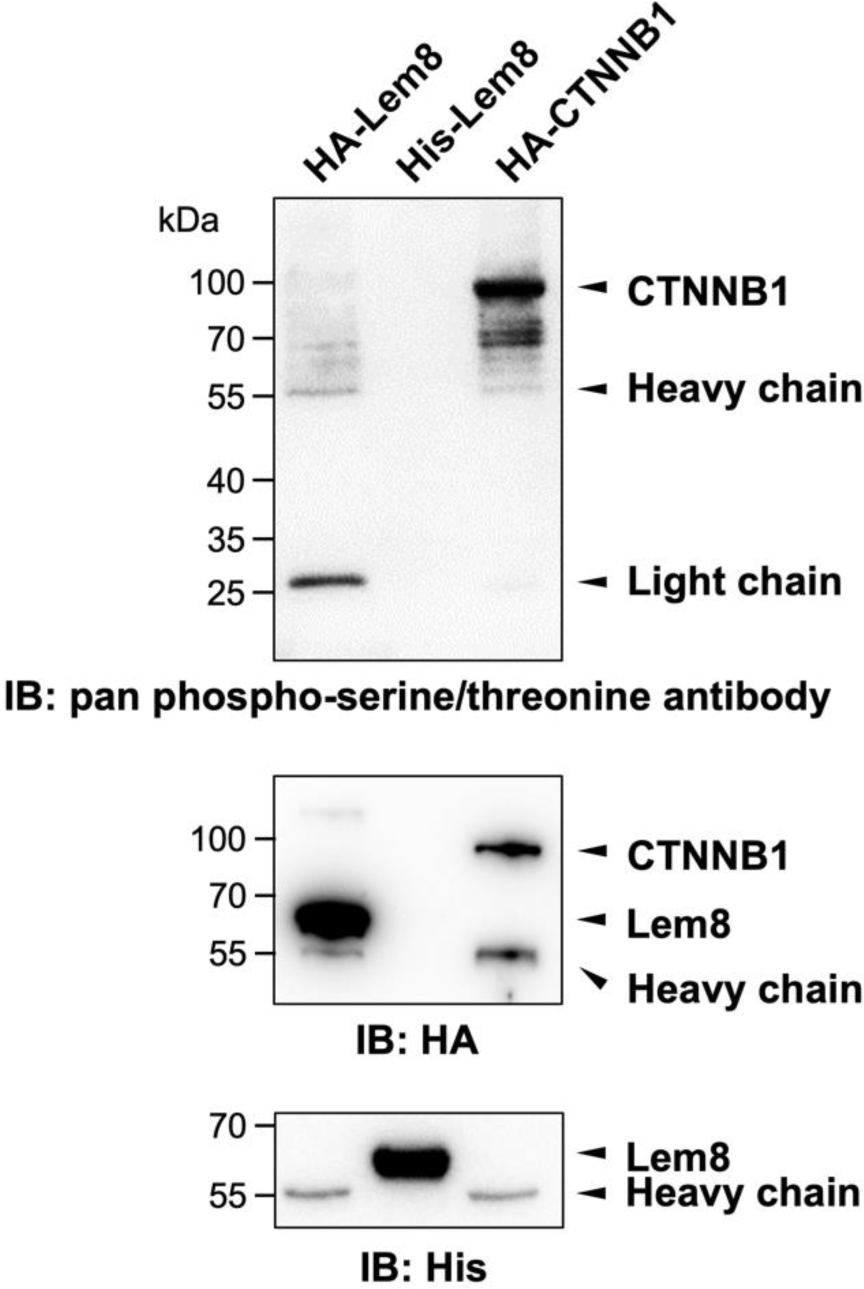
Phospohorylation of Lem8 is not required for 14-3-3ζ binding. Lysates of HEK293T cells expressing indicated HA tagged proteins were subjected to immunoprecipitation with agarose beads coated with the HA antibody. The precipitates, as well as His_6_-Lem8 purified from *E coli* were resolved by SDS-PAGE and probed by immunoblotting with a pan phospho-serine/threonine antibody, the HA specific antibody and the His_6_-specific antibody, respectively.

**Fig. S4.**
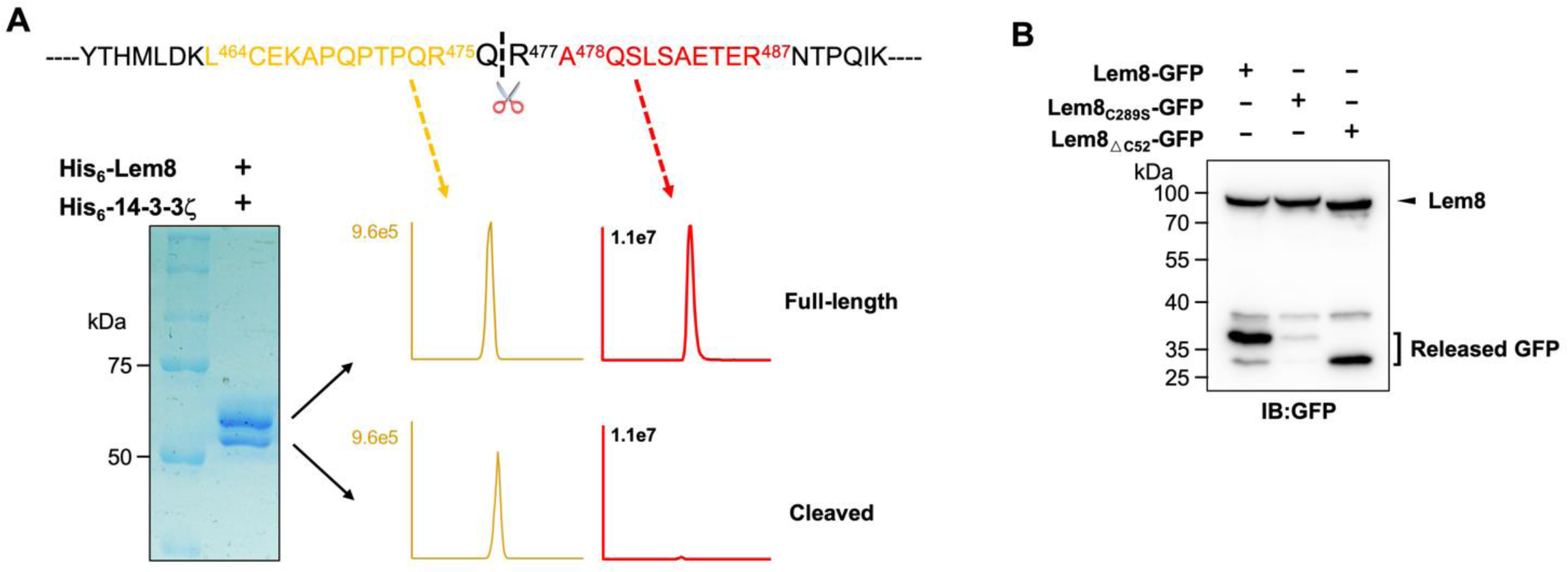
Identification of the self-cleavage site of Lem8. **A.** Determination of the self-cleavage site of Lem8 by mass spectrometry. A diagram of the sequence containing the recognition site with the two diagnostic peptides used to determine the cleavage site (top panel). Protein bands corresponding to full-length and cleaved Lem8 band was excised ( lower left panel), digested with trypsin and analyzed by mass spectrometry. The semi-tryptic peptide -L_464_CEKAPQPTPQRQ_476_- is present in cleaved samples but not in samples of full-length Lem8, whereas the fragment - A_478_QSLSAETER_487_- was only detected in samples of the full-length protein (lower right panel), supporting the notion that the cleavage site lies between Gln476 and Arg477 described in Fig.3C. **B.** Self-cleavage of Lem8 removes GFP fused to its carboxyl end. GFP was fused to the indicated alleles of Lem8 and the fusion proteins were individually expressed in HEK293T cells by transfection. Samples resolved by SDS-PAGE were detected by immunoblotting with GFP-specific antibodies. Results shown were one representative from three independent experiments with similar results.

**Fig. S5.**
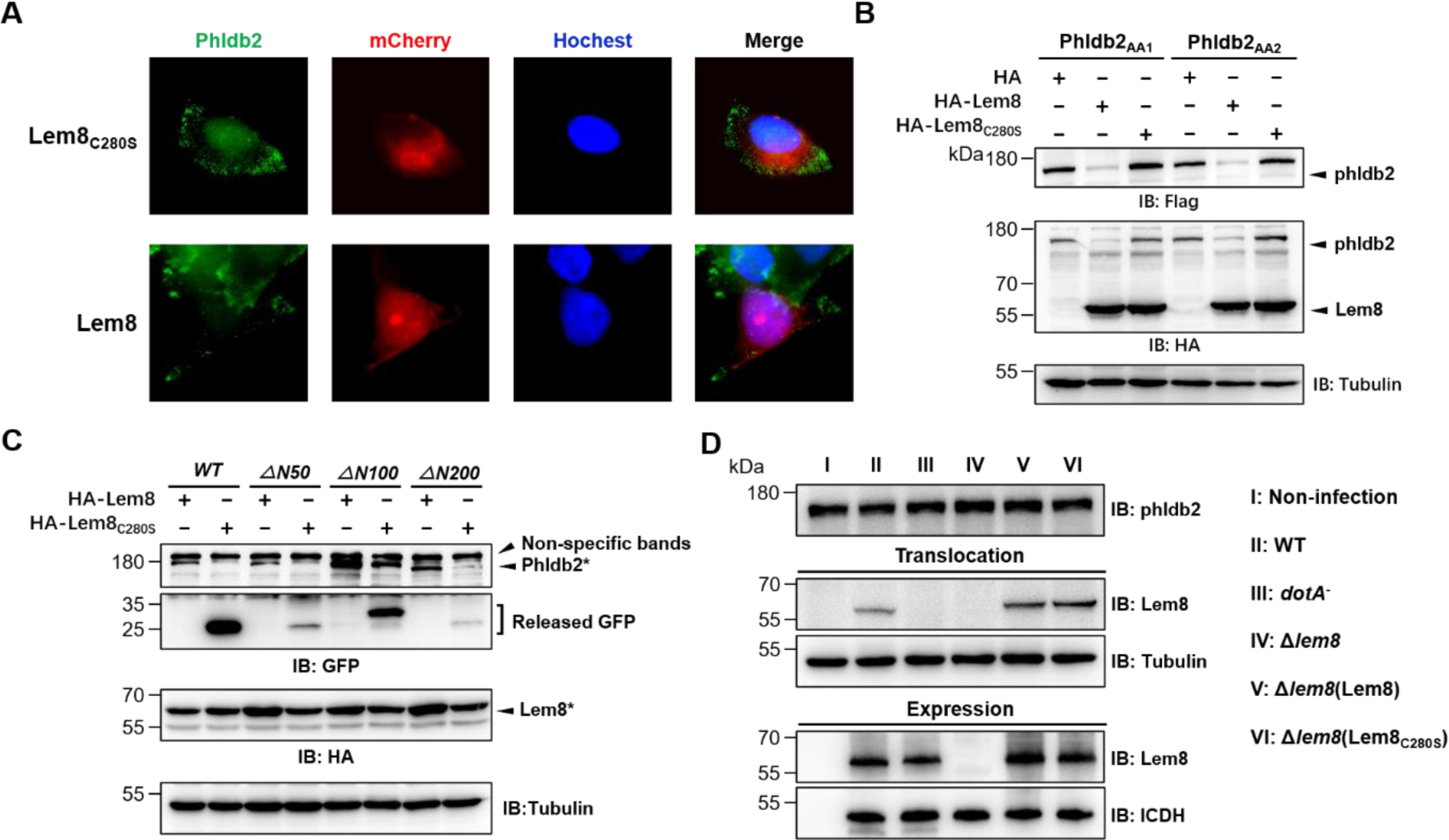
Lem8 cleaves phldb2 at multiple sites. **A.** Lem8 causes redistribution of Phldb2 in cells. Hela cells were transfected to express the indicated mCherry fusion proteins. 24 h after transfection, cells were fixed and immnostained with anti-Phldb2 antibodies. The nuclei were stained by Hoechst 33342. Images were acquired with a Zeiss LSM 880 confocal microscope. Phldb2, green (GFP); Lem8 and its mutants, red (mCherry); nuclei, blue (Hoechst). Bar, 5 μm.Mutations in the cleavage site of Phldb2 cannot completely prevent its degradation by Lem8. **B.** Mutations were introduced into HA-Phldb2-Flag to replace residues Arg_1111_ and Gln_1112_ (Phldb2_AA1_) or Gln_1112_ and Arg_1113_ (Phldb2_AA2_) with alanine, respectively. The two mutants were co-expressed in HEK293T cells with Lem8 or Lem8_C280S_. Samples were resolved with SDS-PAGE, and probed by immunoblotting with the antibody specific to HA and Flag, respectively. **C.** Lem8 removes the GFP tag fused to the amino end of Phldb2 deletion mutants. GFP was fused to the amino end of Phldb2 and the indicated truncation mutants. The fusion proteins were individually co-expressed with HA-Lem8 or HA-Lem8_C280S_ in HEK293T cells by transfection. Samples resolved by SDS-PAGE were detected by immunoblotting with GFP-specific antibodies. Results shown were one representative from three independent experiments with similar results Mutations in the cleavage site of Phldb2 cannot completely prevent its degradation by Lem8. **D.** Cleavage of Phldb2 is undetectable during *L. pneumophila* infection. HEK293T cells transfected to express FcγRII receptor were infected with the indicated bacterial strains. 2 h after infection, the protein levels of Phldb2, as well as the translocation and expression of Lem8, were probed with the appropriate antibodies with Tubulin and ICDH as loading control, respectively. Bacterial strains: I, Non-infection; II, Lp02 (WT); III, *dotA*_-_ (defective in Dot/Icm); IV, Lp02Δlem8; V, Lp02Δlem8(pLem8); VI, Lp02Δlem8(pLem8_C280S_).

**Table S1.**
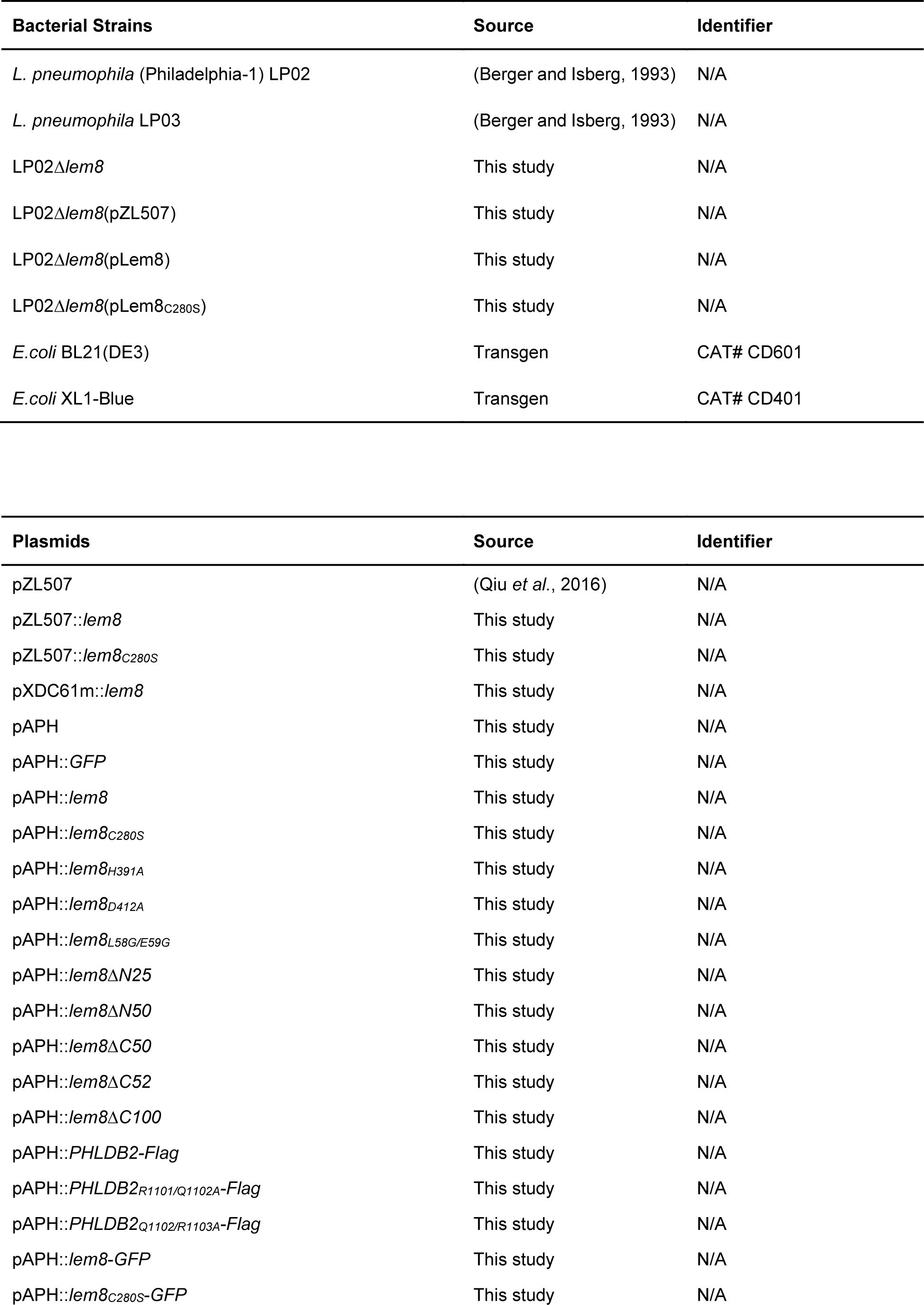
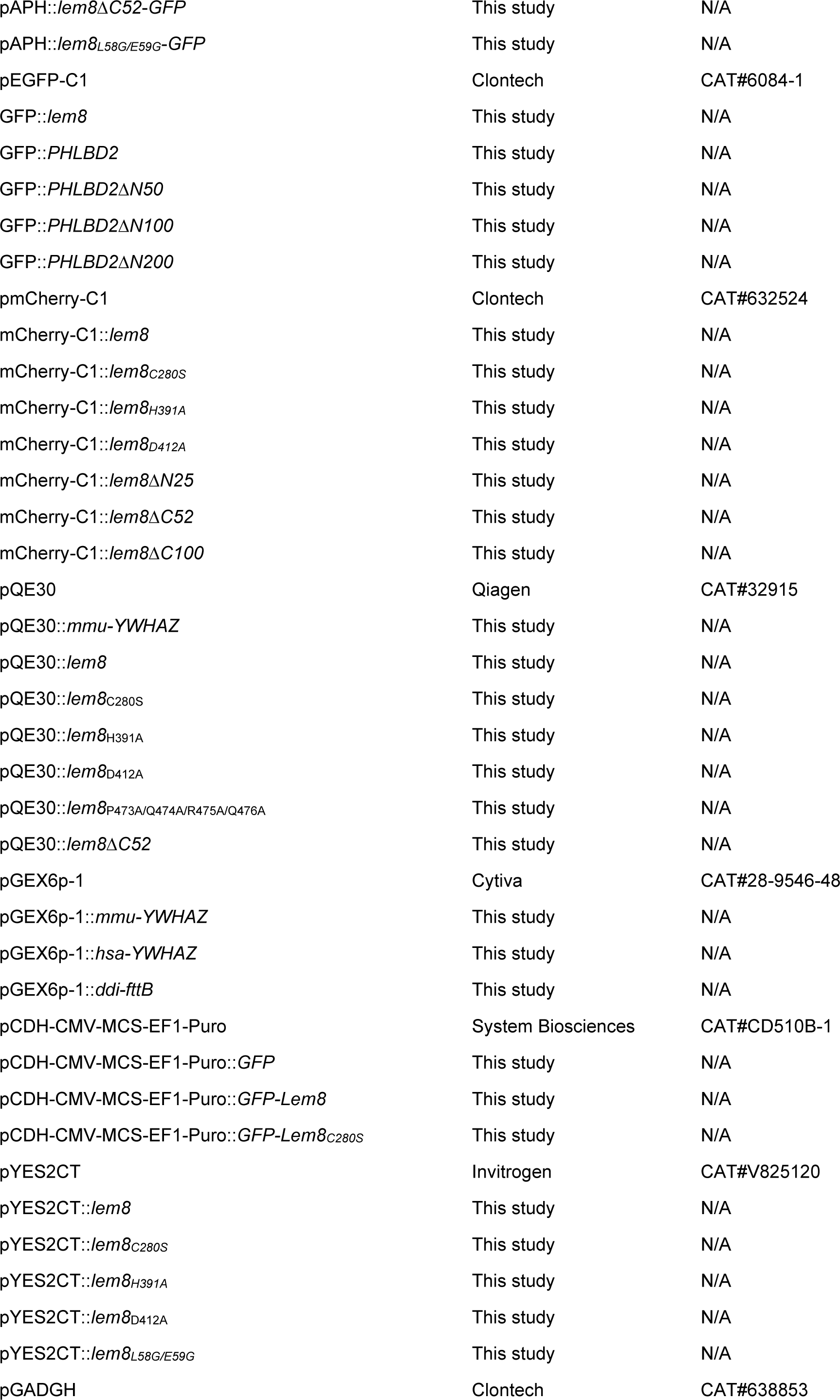
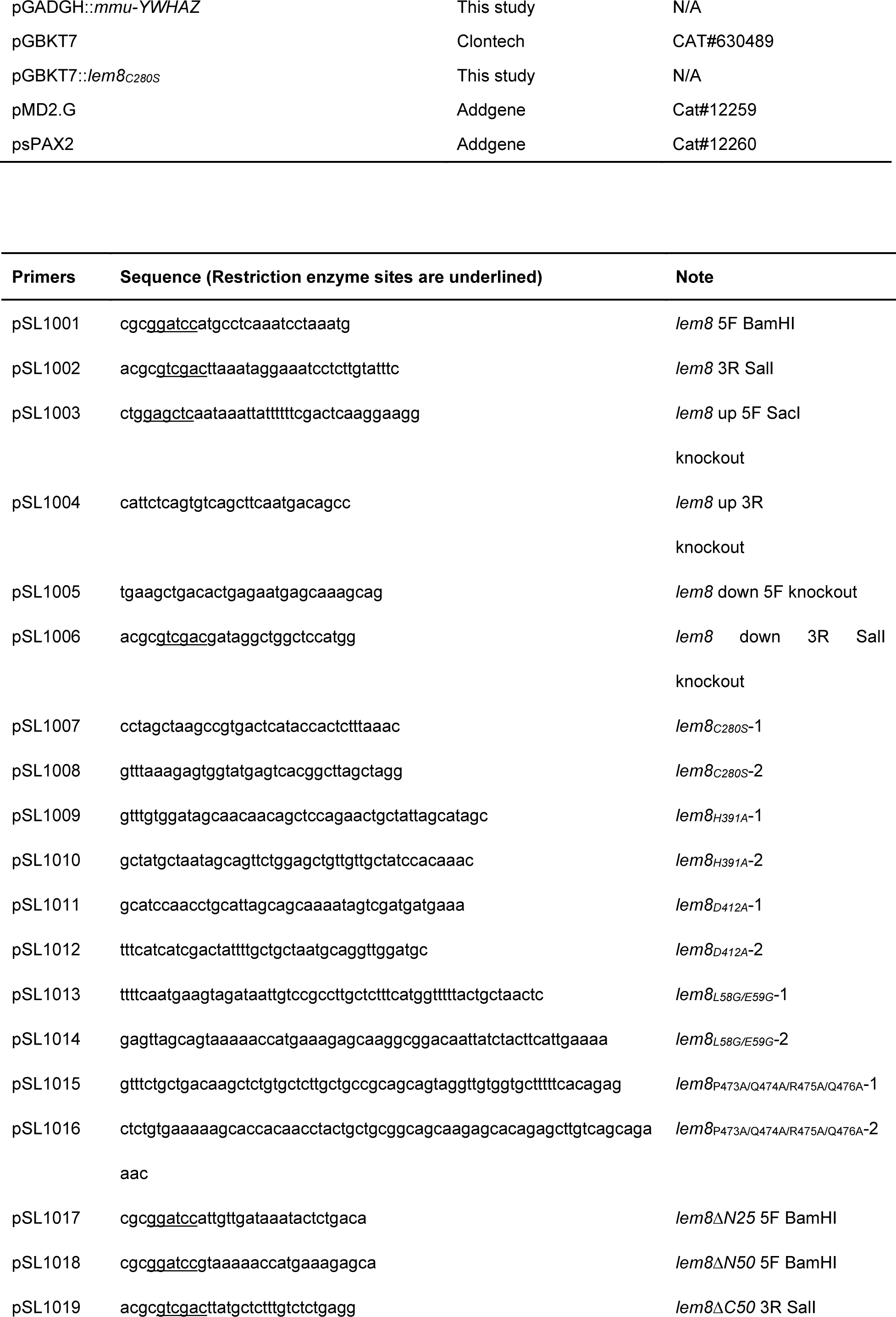
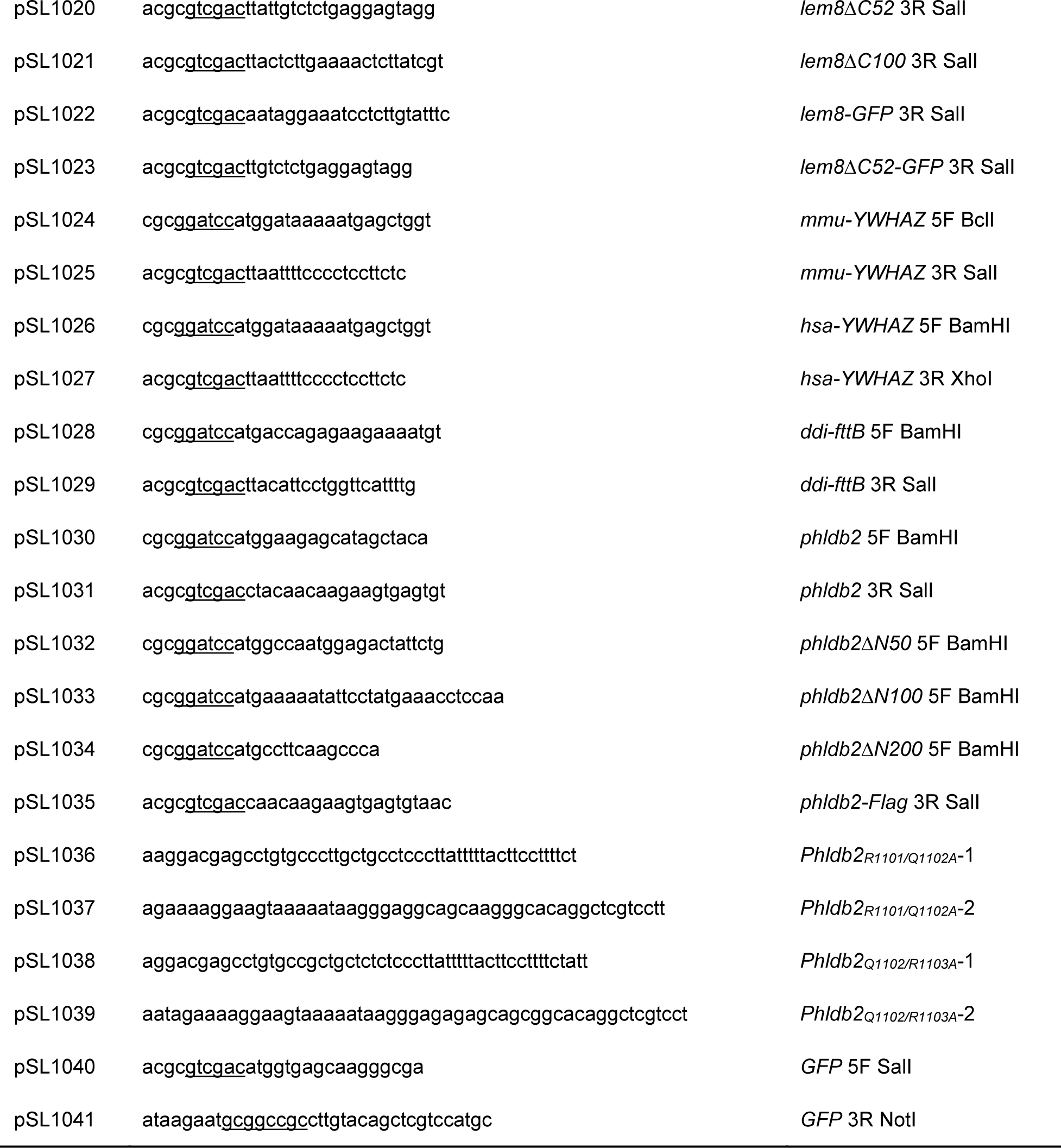
Bacterial strains, plasmids and primers used in the study.

